# Species identity and composition shape productivity of stony corals

**DOI:** 10.1101/2024.06.20.599878

**Authors:** Jana Vetter, Jessica Reichert, André Dietzmann, Lisa Hahn, Anna E. Lang, Giulia Puntin, Maren Ziegler

## Abstract

Coral biodiversity has an enhancing but saturating effect on community productivity, however, the direct effects of neighbouring coral colonies on productivity remain poorly understood due to the complex interplay of biotic and abiotic factors. We set up a fully controlled aquarium experiment, in which we quantified the effects of species identity and composition on the productivity of nine stony coral species from three families. Baseline productivity and the response to neighbouring organisms strongly differed between species. Regardless of whether species increased or decreased productivity, the responses were consistently more pronounced and positive towards conspecific than heterospecific neighbours, indicating kin selection effects between closely related species. Species productivity in monoculture and productivity in polyculture were inversely correlated, with inherently less productive species overperforming in polyculture and vice versa. Our results highlight that contact-free interactions in marine animals shape biodiversity-productivity effects otherwise known from plant communities.

## Introduction

Coral reefs are the most biodiverse marine ecosystems, making them productivity hotspots of ecological and economical significance (Reaka-Kudla, 1997). However, the link between the biodiversity and the productivity of these iconic ecosystems is poorly understood. This lack of information is especially concerning given that coral reefs are particularly affected by climate change (Hughes et al., 2018) and a further loss of up to 90 % of reef ecosystems is predicted even under optimistic climate change projections for the year 2100 (IPCC 2018).

Biodiversity and productivity of ecosystems have complex links and feedback loops. Productivity shapes species richness, and species richness in turn influences ecosystem productivity (Cardinale et al., 2007; Duffy et al., 2017; Mittelbach et al., 2001). The productivity of the species in a community is determined by resource partitioning and niche complementarity (Cardinale et al., 2007; Loreau & Hector, 2001), but also by direct interactions with neighbouring organisms (Engelhardt et al., 2023; Sharifi & Ryu, 2021). For terrestrial plants, these direct interactions between neighbours that are mediated without physical touch either enhance or reduce their productivity (Gorzelak et al., 2015; Sharifi & Ryu, 2021). While heterospecific interactions among plants are mostly dominated by competition, conspecific neighbours mostly facilitate each other (Sharifi & Ryu, 2021). This is in line with the kin selection theory, which posits that genetically close organisms have an evolutionary incentive to help each other (Birch & Okasha, 2015). In addition, baseline productivity of a species may affect its productivity in polyculture, with a so-called negative selection effect occurring when inherently less productive species perform better in polyculture than expected (Hector et al., 2002). Associated trade-offs can result in different interaction strategies depending on the identity of the neighbours, such as e.g., reorienting leaf growth to avoid shading of kin neighbours (Crepy & Casal, 2015) or switching between ‘confrontational’ or ‘avoidance’ growth, depending on the likelihood of successful competition (Gruntman et al., 2017).

Although studies historically focused on terrestrial ecosystems, links between biodiversity and productivity have also been documented in aquatic environments (Bruno et al., 2005; Cardinale et al., 2009; Clements & Hay, 2019, 2021). Similar interactions as in terrestrial plants may potentially occur between stony corals, as the seawater surrounding coral colonies carries their unique microbial and biochemical signature and corals may distinguish self from non-self (Ochsenkühn et al., 2018; Rinkevich, 2004; Tout et al., 2014; Weber et al., 2019). Indeed, conspecific coral aggregations often exhibit increased productivity in the field (Dizon & Yap, 2005; Raymundo, 2001; Shantz et al., 2011). Furthermore, field experiments in the reef have shown primarily positive but saturating effects of coral species richness on growth rate (Clements & Hay, 2019, 2021; Dizon & Yap, 2005), indicating the presence of a link between biodiversity and productivity in stony coral communities. These *in situ* experiments give first insights into the complex biodiversity-productivity relationships of stony corals, yet they cannot reliably distinguish between the many biotic and abiotic factors present in the reef that may drive these relationships.

This study aimed to quantify the productivity of stony corals in response to neighbouring coral species assemblages. We hypothesized that corals physiologically respond to the presence of neighbouring colonies with the identity and composition of the tested coral and its neighbouring organisms determining the direction and strength of the response. To specifically test the kin selection theory and negative selection effect, we set up a fully controlled experiment that excluded confounding factors and quantified contact-free effects of species diversity and composition on the productivity of nine stony coral species from three families (Acroporidae, Poritidae, and Pocilloporidae). Our main objectives were i) to assess the baseline productivity of the corals in monoculture, ii) to quantify productivity changes in polyculture assemblages of coral neighbours from the same species, the same family, or different families, and iii) to determine the relationship between species productivity in monoculture and its productivity in polyculture.

## Methods

### Coral husbandry

Nine coral species from three families were used in the experiment, including Acroporidae: *Acropora cytherea* (Dana, 1846), *Acropora muricata* (Linnaeus, 1758), *Montipora digitata* (Dana, 1846); Poritidae: *Porites lobata* (Dana, 1846), *Porites cylindrica* (Dana, 1846), *Porites rus* (Forskål, 1775); and Pocilloporidae: *Pocillopora verrucosa* (Ellis & Solander, 1786), *Pocillopora damicornis* (Linnaeus, 1758), *Stylophora pistillata* (Esper, 1792). This species selection represents a wide range of morphological and functional groups (Madin et al., 2016). All coral species have been cultured in the Ocean2100 aquarium facility of Justus Liebig University Giessen, Germany, for several years (Tab. S1). For each species three coral colonies were each fragmented into three pieces of 3-5 cm length, resulting in nine coral fragments per species. Fragments were suspended from nylon lines and held in 265-liter tanks, separated by species for at least 3 weeks before the measurement period. This ensured a naïve state of the corals without direct exposure to stimuli from neighbouring organisms. The tanks were part of a 7,000-liter artificial seawater recirculation system, containing additional scleractinian corals, as well as other reef organisms. All corals were fed daily with frozen copepods, at least 2 h before the incubations. Corals were exposed to a 10:14 h light:dark cycle at ∼200 µmol photons m^-2^ s^-1^.

### Experimental design

Coral productivity was measured in 1-liter incubations (light and dark) over a total measurement period of 21 days. Four different types of incubations were performed (Fig. 1; 1,020 incubations in total): i) monoculture incubations, in which each fragment was incubated alone (3 genotypes per species, n = 3 per genotype, n = 9 per species). These single-fragment incubations were conducted during the first, middle, and final three days of the measurement period. For direct comparison the same coral fragments were used in three types of polyculture incubations consisting of multiple fragments. ii) Conspecific incubations, consisting of three coral fragments from three genotypes of one species (9 species, n = 6 per species). iii) Single-family incubations, consisting of three coral fragments from three species of one coral family (3 families, n = 9 per family). iv) Multi-family incubations, consisting of three coral fragments from species of three different coral families (n = 9 per mixture). For the multi-family incubations, 12 species combinations were randomly chosen out of the 27 possible combinations. A species richness of three per polyculture incubation was chosen, based on the findings of Clements and Hay (2021), who identified a saturating effect of species richness of three in their biodiversity-productivity field study, which included six of our nine study species.

**Figure 1.**
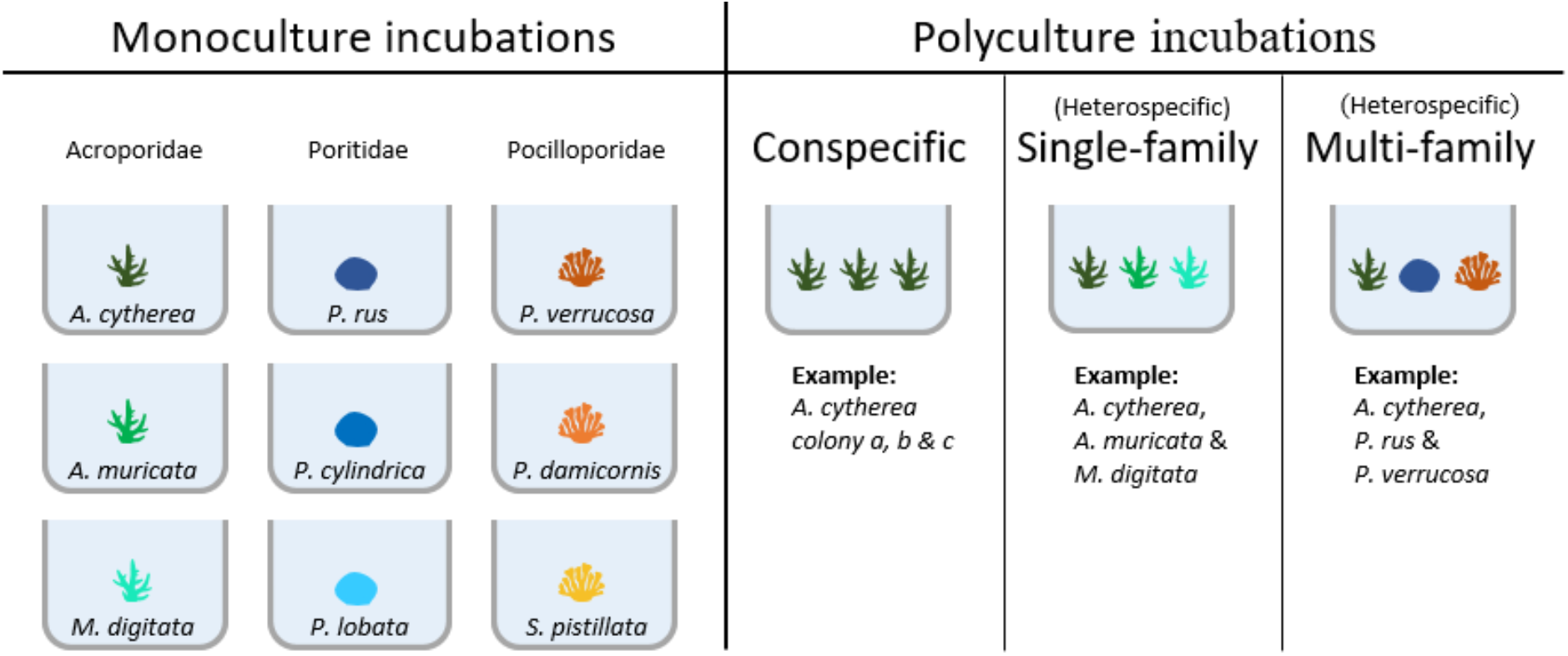
Overview of productivity measurements with different genotype and species combinations. Stony coral productivity measurements from the monoculture (single-fragment) incubations were used to investigate productivity changes caused by neighbouring coral fragments in the three polyculture (multi-fragment) incubation types: conspecific, single-family, and multi-family.

As it is known that coral fragments of various sizes can have differing productivity (Nielsen et al., 2022; Vollmer & Edmunds, 2000), the fragments in this study were all within a size range (12-44 cm^2^), which was above the critical minimal size previously identified by Vollmer & Edmunds (2000). The fragment sizes differed per species with 15.8 ± 3.0 cm^2^ to 31.6 ± 7.5 cm^2^ (mean ± SD). To check for biomass effects, we additionally conducted incubations, with three fragments of the same coral genotype (n = 3, Tab. S2). Coral density, hence a biomass increase, has been shown to affect coral productivity (Ladd et al., 2016; Shantz et al., 2011), yet we found no such correlation except for net photosynthesis and calcification of *P. verrucosa* (Tab. S2).

Per day, three incubation runs, each consisting of a 30-minute light and a 30-minute dark incubation, were performed. The timeframe was chosen based on values from pilot experiments, to not exceed critical oxygen over- or under-saturation. The first incubation began approximately three hours after the lights in the holding tanks were turned on and every 2.5 h from then (i.e., 10:00 h, 12:30 h, and 15:00 h). Each coral fragment was only used once per day and alternated between runs, to control for daytime effects. This resulted in a fully crossed design. Incubations took place in airtight, 1-liter glass jars, mimicking conditions in the holding tanks at constant temperature (26.5 °C), flow rate (6 cm s^-1^), salinity (35 ppt), and light conditions (168 ± 2 µmol photons m^-2^ s^-1^), using a multi-point stirring incubator (Rades et al. 2022). Seawater from an empty tank of the recirculation system was used for the incubations and fragments were positioned so that they did not touch the glass or each other at an approximate distance of 5 cm between fragments. Each incubation run included nine coral incubation jars and one seawater control jar without organisms to account for microbial activity in the water.

### Productivity parameters

#### Photosynthesis and respiration

Net photosynthesis and respiration were derived from light and dark incubations, respectively. Changes in dissolved oxygen (DO) were measured based on the difference in DO concentration before and after each incubation with an optical oxygen multiprobe (FDO 925, WTW Multi 3620 IDS, Weilheim, Germany). DO changes were corrected with respective seawater controls, normalized to incubation time, water volume, and coral surface area. Gross photosynthesis was calculated as the sum of net photosynthesis and respiration values.

#### Calcification measurements

Coral calcification rate was calculated using the alkalinity anomaly technique (Chisholm & Gattuso, 1991; Gazeau et al., 2015). Corals were held at a constant total alkalinity (TA) of 2,618 ± 33 µmol L^-1^ (mean ± SD). Water samples (30 ml) were taken at the beginning and at the end of each light incubation, stored at room temperature in the dark, and analyzed within 5 h of collection. Total alkalinity was determined using an automated open-cell potentiometric titrator (TitroLine® 7000, SI Analytics, Germany) with a pH and temperature glass electrode (Pt1000, SI Analytics™, Germany), which was calibrated daily (slope from 98 to 99 %). Measurements were carried out at room temperature (20 °C). TA was calculated via the Gran Approximation as per Dickson et al. (2007) and using a modified version of the ‘alkalinity’ function of the seacarb package (Gattuso et al. 2023). Calcification rate (G) was then calculated using the following equation (Schneider & Erez, 2006):

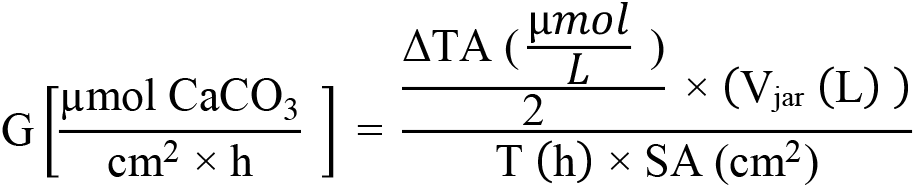

where ΔTA is the difference between TA of the control incubations and TA of the coral incubations at the end of each incubation, V_jar_ is the water volume of the incubation jar (L), T is the incubation time (h), and SA represents the surface area of the incubated coral fragments (cm^2^). The difference between the three controls per day was better than ± 1.6 %, with an average difference of ± 0.5 %.

#### Surface area determination

Photosynthesis, respiration, and calcification rates were normalized to the surface area of each coral fragment. Surface areas were measured at the beginning of the three-week experiment, following a two days rest period, before the first incubation took place. The surface areas were determined from 3D models using a handheld 3D scanner (Artec Spider 3D, Artec 3D, Luxembourg), together with the software Artec Studio 9.2.3.15 (Artec 3D, Luxembourg). Surface area and volume were calculated in MeshLab 2016.12 following Reichert et al. (2016).

### Statistical analysis

All data exploration and statistical analysis took place in R version 4.3.1 (R Core Team, 2023), using the tidyverse collection (Wickham et al., 2019). Two runs of the single-fragment incubations (runs 2 and 3 on day 17) were removed from the analysis, due to incorrectly recorded DO_start_ values. Additionally, twelve other outliers in oxygen values as well as twelve calcification outliers were removed from further analysis (Tab. S2, S4–S6).

Differences in productivity between coral families based on species monocultures in single-fragment incubations were tested with a one-way ANOVA, using nlme package (Pinheiro et al., 2020) followed by Tukey’s post hoc test with Bonferroni correction, using the multcomp package (Hothorn et al., 2008). To account for the multiple measurements of the same fragments, ‘fragment ID’ was used as random factor. The response variable ‘gross photosynthesis’ was square root transformed to meet test assumptions.

To establish the effect of species assemblage on productivity, the difference between the measured productivity of a polyculture assemblage and the expected productivity of this assemblage based on the sums of the productivity of the respective monocultures was calculated. To calculate the expected productivity values per polyculture, all values of the monoculture productivity measurements from the three fragments of the same genotype were interpolated for each day and run (Fig. S1). To test for differences between expected and measured productivity a paired t-test with Bonferroni correction was used, as the experimental design allowed for direct comparisons of expected and measured productivity values. This was visualized as the percentage change from expected productivity:

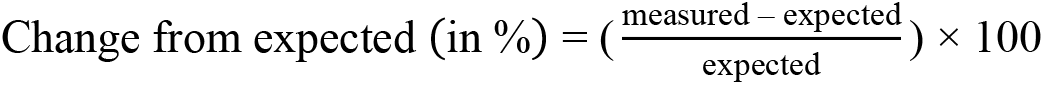

To assess the difference in effect strength between conspecific and multi-family incubations, a Wilcoxon rank-sum test was used on the absolute values of the percentage change from expected. To test the strength of the linear correlation between the change from the expected productivity (measured – expected) and the expected productivity a Pearson correlation was used. As the correlation between expected value and change from expected is inherently negatively biased (Blomqvist, 1977; Chiolero et al., 2013), we adjusted the correlation coefficient (R) according to Blomqvist (1977). Subsequently, the corrected p-value was obtained via Fisher’s z-transformation. To further determine to which degree the performance of species in monoculture explained the performance of species assemblages in polyculture a linear regression model was applied.

It could not be tested whether oxygen levels at the start (DO_start_) and at the end (DO_end_) of incubations or the surface area of fragments within the incubations were additional explanatory factors in this correlation analysis, as the dependent variables are mathematically linked to these parameters (Archie Jr, 1981). For example:

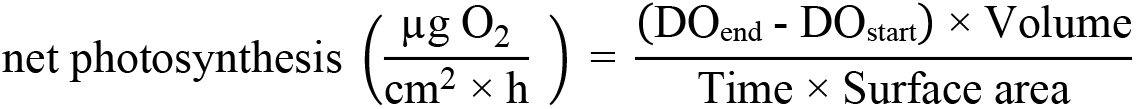

A Pearson correlation between related variables, such as change from expected net photosynthesis and initial DO, where one is derived from the other, cannot be directly calculated due to the same inherent negative bias mentioned above (Blomqvist, 1977; Chiolero et al., 2013). Due to the different units, a correction following the Blomqvist formular is not possible. Therefore, to limit the influence of surface area and DO_start_ values the variables were kept in a similar range over all incubations. A positive correlation with surface area was confirmed for all productivity parameters before normalization with surface area (Fig. S2), indicating a continuous relationship of productivity across fragment size ranges and single vs. multi-fragment incubations. DO_start_ values of the light incubations were kept constant at 7.0 ± 0.1 mg liter^-1^ (± SD). The corals were left in the incubation jars between light and dark incubations to decrease handling stress. Hence corals with a higher net photosynthesis had a higher DO_start_ value for the dark incubations. These incubations tended to have a higher respiration, as expected because highly productive species in the light generally also respire more DO in the dark (McCloskey’ et al., 1978).

## Results

### Productivity differs between stony coral families

Overall, the Pocilloporidae exhibited significantly higher net photosynthesis, respiration, gross photosynthesis, and calcification rates (p < 0.05), than the Acroporidae and Poritidae, which were not significantly different from each other (Fig. 2; Tab. S3-S4). These patterns were based on 27 monoculture incubations for each of the three species within each family. Generally, species from the Pocilloporidae were among the most productive, while species from the Poritidae were among the least productive (Fig. 2). Species from the Acroporidae had the largest range with *M. digitata* among the most productive species and *A. cytherea* and *A. muricata* with intermediate to low productivity (Fig. 2).

**Figure 2.**
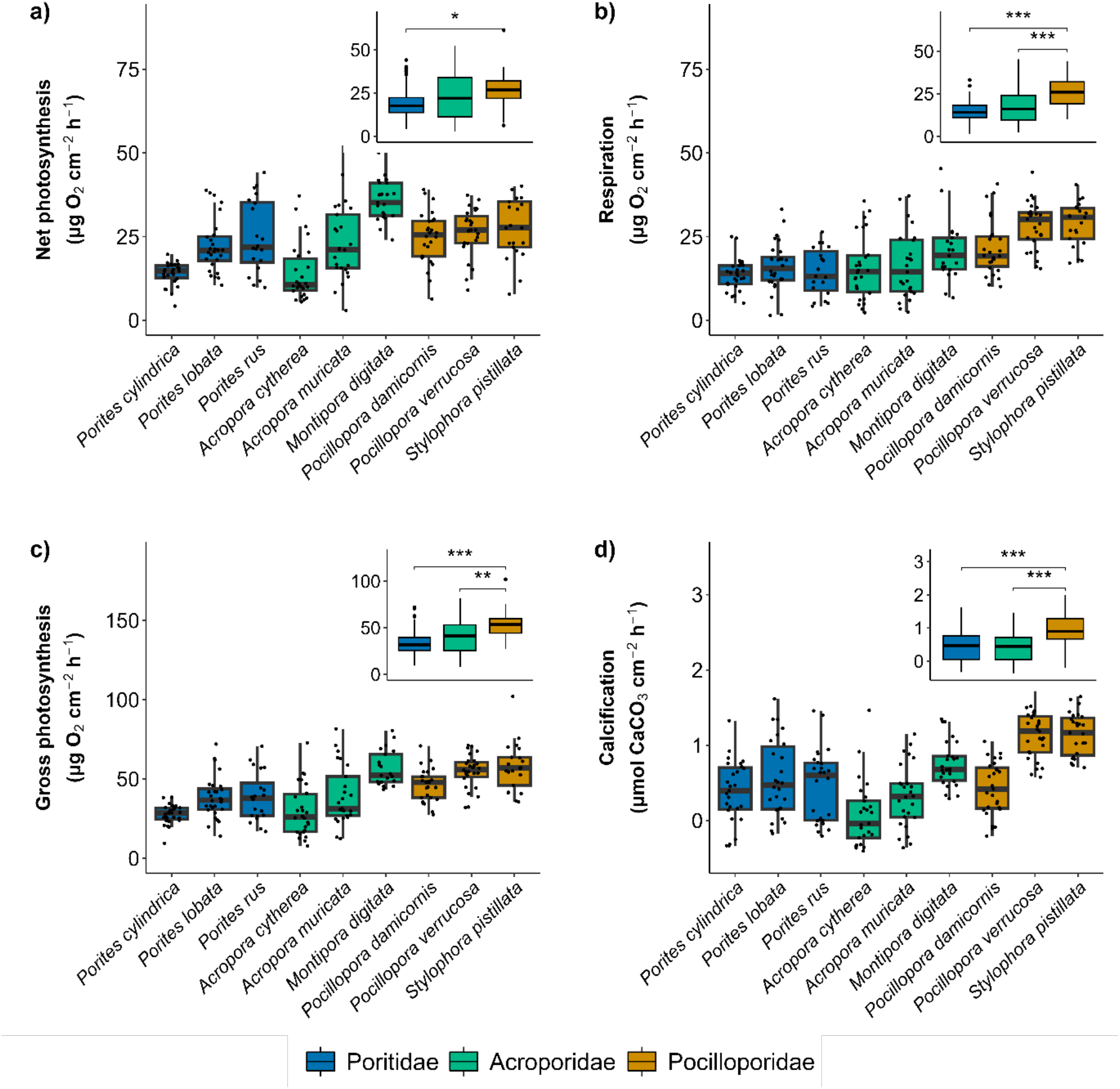
Productivity of nine stony coral species from three families. Productivity was measured as net photosynthesis (a), respiration (b), gross photosynthesis (c), and calcification (d). Data based on monoculture incubations (n = 27). Boxplots are ordered by family, boxes indicate the first and third quartiles, and whiskers indicate ± 1.5 IQR. Asterisks indicate significant differences between families (* p < 0.05, ** p < 0.01, *** p < 0.001).

### Neighbour composition affects stony coral productivity

To test effects of species composition on coral productivity, polycultures (conspecific, single-family, and multi-family) were compared with monocultures of the same species. The presence of conspecific neighbours primarily led to an upregulation in photosynthesis compared to when genotypes were incubated alone (Fig. 3a-c, Tab. S5). These positive effects of genotypic diversity on productivity within species were especially pronounced within the Poritidae family, while the Pocilloporidae family showed least effects. *P. rus* and *M. digiata* were the exception with lower photosynthesis than expected in response to genoptypic diversity. A mix of species within coral families (single-family incubations) had no impact on photosynthesis of the assemblage (Tab. S6), while the presence of more distantly related species from different families (multi-family incubations) often led to a decrease in photosynthesis, and positive effects on productivity were rare (Fig. 3d-f, Tab. S7). Two of these multi-family assemblages (*P. verrucosa* & *M. digitata* & *P. rus*; *S. pistillata* & *M. digitata* & *P. rus*) had lower net photosynthesis than *P. rus*, their least productive species in monoculture. Yet, none of the multi-family assemblages were more productive than their most productive species in monoculture. Calcification was significantly increased in one conspecific, one single-family, and one multi-family assemblage each, but was otherwise largely in the range of the expected values (Tab. S5-S7).

**Figure 3.**
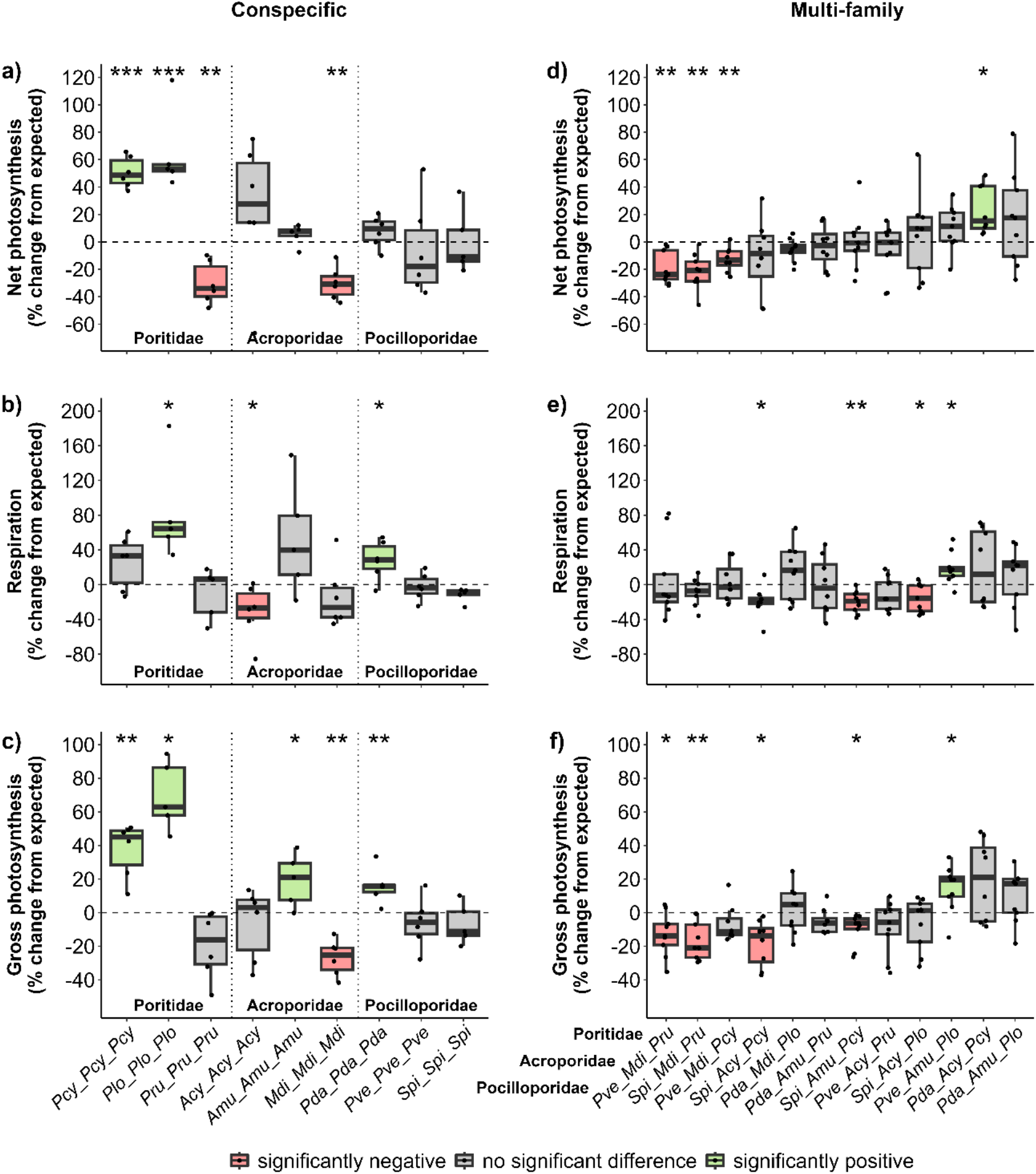
Coral neighbour effects on productivity in response to conspecific (a-c) and multi-family assemblages (d-f). Effects are shown as change from expected productivity (in %) of net photosynthesis (a & d), respiration (b & e), and gross photosynthesis (c & f). Species abbreviations: Poritidae: Pcy = *P. cylindrica*, Plo = *P. lobata*, Pru = *P. rus*; Acroporidae: Acy = *A. cytherea*, Amu= *A. muricata*, Mdi = *M. digitata*; Pocilloporidae: Pda = *P. damicornis*, Pve = *P. verrucosa*, Spi = *S. pistillata*. Red boxes indicate significantly lower, green significantly higher, and grey similar productivity of the assemblage in comparison to the productivity of the same species in monoculture (* p < 0.05, ** p < 0.01, *** p < 0.001). The boxes mark the first and third quartiles, and the whiskers indicate ± 1.5 IQR.

### Coral species have consistent responses to diverse neighbours

Overall, conspecific neighbours triggered three-fold larger changes in photosynthesis than those observed in multi-family assemblages (p < 0.05, Tab. S8, Fig. 4c). For example, the gross photosynthesis in conspecific incubations diverged from expected values by 96 % (-27 to 69 %), while the range in the multi-family assemblages was 37 % (-18 to 19 %). Regardless of whether the presence of conspecifics boosted or inhibited net photosynthesis, respiration, and gross photosynthesis, the response direction was largely consistent between parameters within species (Fig. 4a). The same holds true for the effect in multi-family assemblages (Fig. 4b). Certain species were consistently present in overperforming assemblages, while others were present in underperforming assemblages. Specifically, the species *P. lobata*, *P. damicornis*, and *A. muricata* were mostly present in assemblages that demonstrated higher than expected productivity. In contrast, the species *M. digitata*, *P. verrucosa*, *S. pistillata*, *P. cylindrica*, and *P. rus* were primarily found in assemblages that showed lower than expected productivity. Interestingly, the same species which were found in the overperforming multi-family assemblages also overperformed in the conspecific assemblages, and the species which were found in the underperforming multi-family assemblages also underperformed in the conspecific incubations (Fig. 4). The only exception was *P. cylindrica,* which increased its productivity with conspecific neighbours despite being predominantly present in underperforming multi-family incubations.

**Figure 4.**
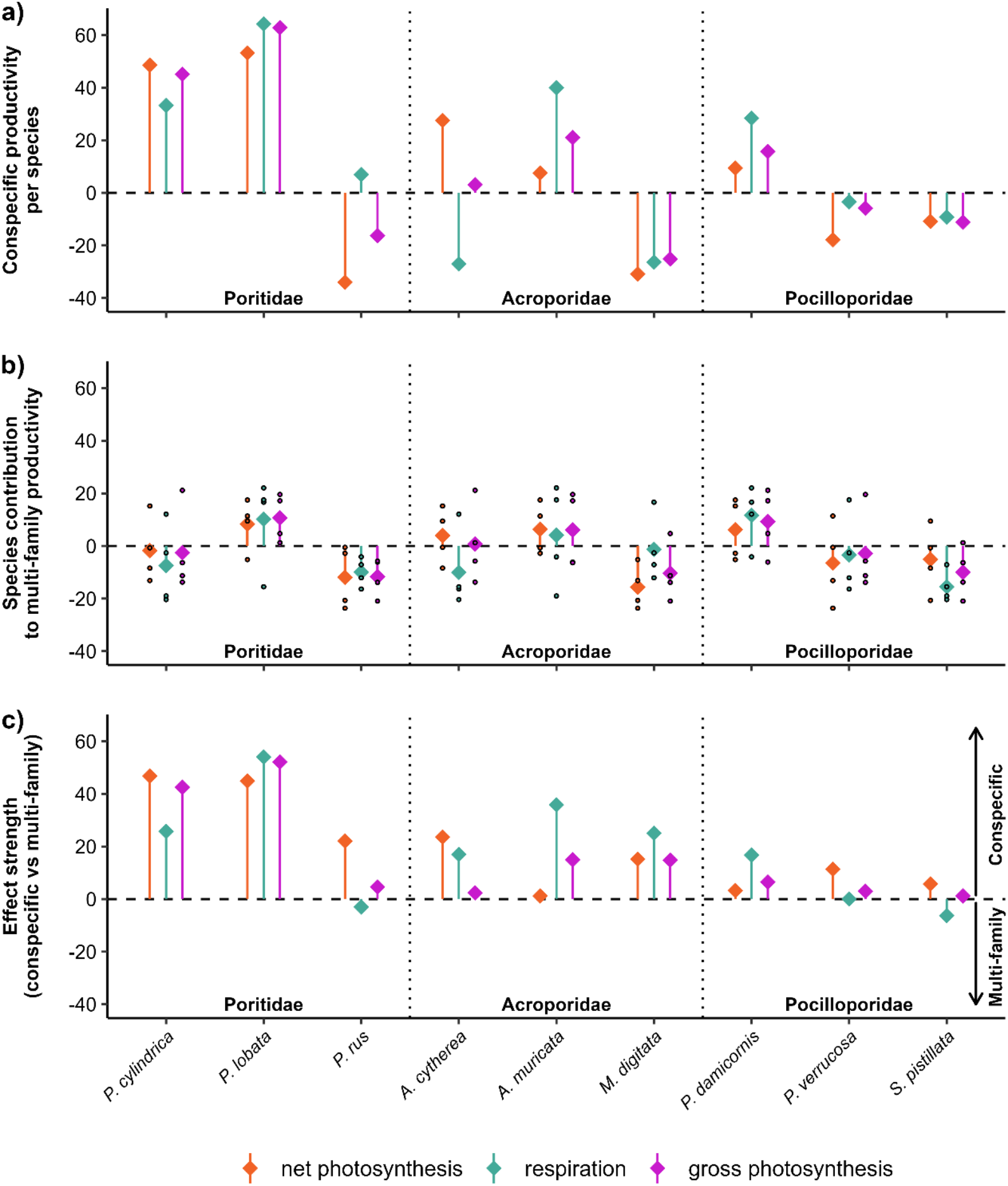
Physiological response to coral neighbour compositions. Responses are shown as overall productivity change from expected productivity (%), calculated as the median per species in the conspecific assemblages (a) and as the average median of all multi-family assemblages per species (b). Black dots indicate the medians of the multi-family assemblages per species. The effect strength of conspecific and multi-family assemblages is compared via their absolute difference (i.e., |conspecific median| - |multi-family median|), for each species (c). Positive values indicate a stronger effect of conspecifics on species productivity, whereas negative values indicate a stronger effect of species from different families on species productivity.

### Productivity of polycultures is linked to coral performance in monoculture

The expected productivity and the measured productivity of all conspecific and multi-family assemblages were positively correlated (Fig. 5a & c, Fig. S3a-c., Fig. S4a-c, Tab. S9). Coral assemblages consisting of highly productive species in monoculture were also more productive in polyculture compared to those assemblages with less productive species in monoculture. This pattern was more pronounced in the conspecific assemblages, where performance in monoculture explained 9-60 % of the performance in polyculture, compared to 9-25 % in the multi-family assemblages. The expected productivity and the deviation from the expected productivity of these assemblages was negatively correlated for net and gross photosynthesis of the conspecific assemblages (Fig. 5b, Fig. S3d-f, Tab. S10) and net photosynthesis, respiration, and gross photosynthesis of the multi-family assemblages (Fig. 5d, Fig. S4d-f, Tab. S10), even after strict correction for the absolute variance within a colony, which might have led to a slight overestimation of the errors-in-variables bias (Blomqvist, 1977). In other words, species assemblages consisting of highly productive species in monoculture were less productive in polyculture than expected, and assemblages consisting of less productive species in monoculture were more productive in polyculture than expected. This led to a slightly more homogenous productivity among the polycultures with a lower relative range between minimum and maximum values in comparison to monocultures. Again, the conspecific assemblages exhibited a more pronounced pattern than the multi-family assemblages, with a stronger negative correlation between expected and change from expected net and gross photosynthesis (Fig. 5b & d, Tab. S10).

**Figure 5.**
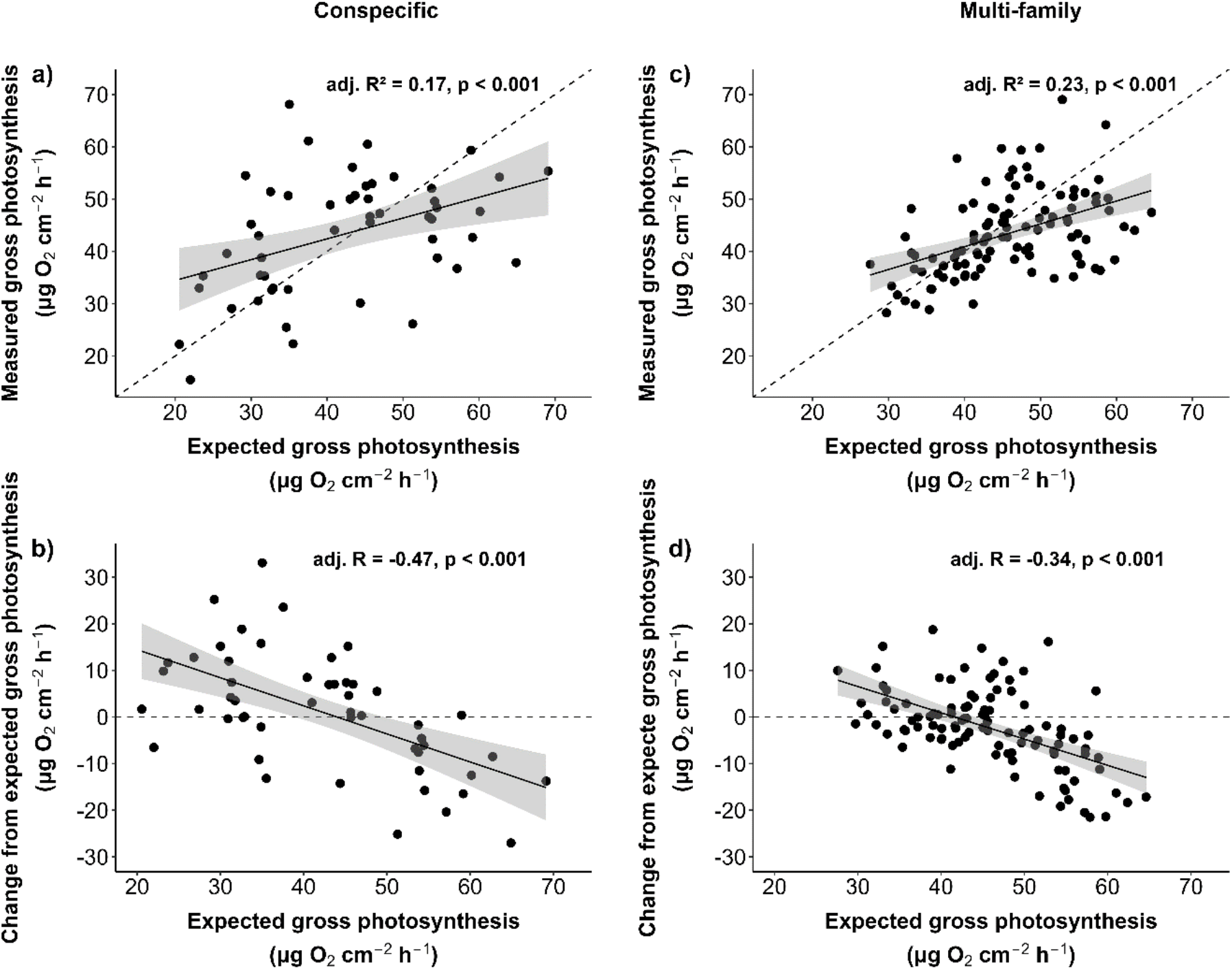
**Relationship between gross photosynthesis of corals in monoculture and polycultures of conspecific (a-b) and multi-family (c-d) assemblages**. Data shown as the relationship between the measured productivity of polyculture assemblages and their expected productivity, displayed with adjusted R^2^ and p-values from a linear regression model (a & c). The productivity change of polyculture assemblages (measured - expected) and their expected productivity (b & d) with corrected R and p-value following Blomqvist (1977). Dashed lines indicate the 1:1 productivity (no change). Points above the dashed lines are assemblages which were more productive in polyculture (conspecific: n = 51; multi-family: n = 104).

## Discussion

Biodiversity-productivity effects in coral reefs may be partially attributed to contact-free interactions between stony corals, as evidenced by our controlled incubation experiment. We found that nine coral species exhibited immediate physiological responses to both conspecific and heterospecific neighbours. Depending on the species composition, productivity was either boosted or reduced, suggesting that corals possess the ability to sense neighbouring corals and modulate their productivity in response. Interestingly, conspecific assemblages had a stronger and generally more positive effect on species productivity than species from other families. The observed changes in productivity from different coral assemblages could explain parts of the biodiversity-productivity effects reported from the field, where functional links are much harder to disentangle due to many confounding factors.

### Baseline productivity of coral families varies

Species in monoculture exhibited family-specific productivity patterns. Specifically, Poritidae demonstrated the lowest photosynthesis and respiration rates, while displaying intermediate calcification rates. This observation corroborates previous findings, indicating substantial allocation of photosynthetic energy into calcification (Carlot et al., 2022). Pocilloporidae had higher photosynthesis and calcification rates than the other two families. *P. verrucosa* and *S. pistillata* even exceeded their typical global calcification spectrum by 200 %, while the other species were within reported ranges (Roik et al., 2016). The high photosynthetic activity in the Pocilloporidae likely also manifested in their high calcification rates (Mallon et al., 2022). The Acroporidae were the only family to show a wide spread in their productivity, with *M. digitata* consistently on the higher end and *A. cytherea* on the lower end of the range. This might reflect their different life history strategies, with the former being considered a weedy and the latter a competitive species (Madin et al., 2016).

### Neighbours regulate coral productivity

We demonstrated that the productivity of stony corals is influenced by the neighbouring coral species and while both up- and downregulation of productivity was observed, conspecific interactions predominantly increased productivity, indicating that the kin selection theory might also apply to corals (Sharifi & Ryu, 2021). In other words corals have an evolutionary incentive to support genetically close neighbours and these facilitative effects between closely related corals might lead to a positive feedback loop (Mittelbach et al., 2001; Shantz et al., 2011). High abundance of conspecific neighbours might result in higher reproductive success (Elahi, 2008) or competitive advantages (Idjadi & Karlson, 2007). This positive effect of conspecifics aligns with previous studies that observed positive productivity effects in the presence of conspecific coral neighbours (Dizon & Yap, 2005; Raymundo, 2001; Shantz et al., 2011), with the potential to mitigate anthropogenic pressures such as ocean acidification (Evensen & Edmunds, 2016).

Various microbes, such as fungi (Roik et al., 2022) and bacteria (Weber et al., 2019), might play an important role in facilitating positive interactions. For example, corals or their associated microbes generate indole-3-acetic acid (IAA), a metabolite that may augment photosynthetic productivity (Weber et al., 2022), by enhancing Symbiodiniaceae growth (Matthews et al., 2023). Coral species such as *P. lobata* which almost doubled its productivity when incubated with conspecifics might either have higher densities of these facilitating microbes or be more receptive to their stimuli. It has previously been shown that corals can react to bacteria within minutes with a release of reactive oxygen species such as hydrogen peroxide (Armoza-Zvuloni et al., 2016), which might trigger a variety of physiological responses, including in photosynthesis and respiration (Quan et al., 2008). It should be noted that conspecific neighbours can also have negative effects on productivity. For example, by harboring host specific pathogens, which have been shown to cause increased mortality of coral larvae in water which surrounds conspecific adults, in comparison with water from around heterospecific corals (Marhaver et al., 2013).

The Poritidae family generally exhibited the strongest response to conspecifics in our study. This corresponds well with a field study in which Poritidae corals grew 30 % more when conspecific density was elevated (Shantz et al., 2011). All Poritidae in our study were gonochoric with each individual either male or female, whereas the species from the other families were hermaphrodites with each individual carrying both sexes (Baird et al., 2009; Madin et al., 2016). Hence, one possible explanation for the strong response in the Poritidae might be that gonochoric species react stronger to conspecific neighbours which might specifically increase their reproductive success.

In contrast to the conspecific assemblages, corals in the multi-family assemblages often exhibited a decrease in productivity. Distinct competitive strategies among the nine studied stony coral species may explain this pattern (Chadwick & Morrow, 2011). Specifically, more aggressive coral species may allocate energy into defense mechanisms, such as mesenterial filaments or sweeper tentacles (Lapid & Chadwick, 2006), when neighbouring corals are present, whereas others may use extra energy for growth as an outcompeting strategy (Rinkevich & Loya, 1983, 1985b; Romano, 1990). However, such strategies could not be directly linked to the observed productivity pattern, as stress-tolerant, weedy, and competitive species were indiscriminately distributed among the top and worst performing species (Madin et al., 2016). The interpretation of the productivity patterns is further hampered by the unknown extent to which the species downregulated their own productivity or negatively influenced the productivity of their neighbours. Two species assemblages that were less productive than their least productive species in monoculture suggests severe hampering of the productivity of all involved species. This contrasts with macroalgae assemblages, where complementarity effects in long-term polyculture were always positive (Bruno et al. 2005). Overall, the response to heterospecific neighbours was not as strong as that to conspecifics, possibly due to a neutralizing effect, which likely occurred in the single-family and multi-family incubations. In other words, if some species react with an increase in productivity while others react with a decrease, the result is a net zero change.

### Species reaction is consistent across neighbours

Interestingly, the response direction in the productivity of each species remained generally unchanged across conspecific and multi-family assemblages, with certain species consistently present in overperforming assemblages and other species in underperforming assemblages. This strikingly similar pattern indicates a predetermined strategy to deal with coral neighbours which might be specific to each coral species. Such a pattern has been observed before and also includes responses to other benthic organisms such as soft corals, algae, and sponges (Engelhardt et al., 2023). For instance, in our study both *M. digitata* and *P. rus*, which demonstrated the strongest productivity decrease with diverse neighbours, also exhibited the strongest productivity decrease to conspecific neighbours. *P. cylindrica* was the only exception to this overarching productivity pattern with downregulated productivity in multi-family assemblages and upregulated productivity in conspecific assemblages. This pattern is similar to previous reports of *P. cylindrica* decreasing productivity among heterospecific coral neighbours (McWilliam et al., 2018), and increasing productivity among conspecifics (Dizon & Yap 2005). Yet, field data suggests that productivity of *P. cylindrica* may also increase in heterospecific assemblages (Clements & Hay, 2019; Dizon & Yap, 2005), which might be explained by long-term competition in the field (Romano, 1990) or other biotic (e.g., predators; Johnston & Miller, 2014) and abiotic effects (e.g. currents; McWilliam et al. 2018). Our study hereby provides a unique dataset, which sets a baseline for further investigations that are needed to disentangle the various parameters affecting biodiversity-productivity interactions in coral reefs.

### Contact-free signalling pathways may mediate productivity changes

Importantly, our results reveal that corals sense the presence of neighbouring coral colonies without direct contact and that this sensory signal results in a productivity change. Corals down- or upregulate their physiological functions as an immediate reaction to the proximity of other coral species in anticipation of competition or facilitation. This behaviour has been observed in terrestrial plants, where a complex web of contact-free interactions exists (Bilas et al., 2021), whereby the type of their physiological response alternated between competing (Gruntman et al., 2017) and facilitating plant neighbours (Crepy & Casal, 2015).

Several signalling pathways potentially mediate the physiological processes: 1) Perception of changes in the light spectrum through reflection from the surface of neighbours. This is the best-known signalling method for the ‘shade avoidance syndrome’ (SAS) (reviewed by Roig-Villanova & Martínez-García, 2016), whereby terrestrial plants sense the proximity of other plant species and react via physiological changes in order to avoid future shading (Bou-Torrent et al., 2014; Gruntman et al., 2017; Roig-Villanova & Martínez-García, 2016). SAS strategies have been proposed for macroalgae and coral, yet have rarely been tested in marine environments (Dizon & Yap, 2005; Monro & Poore, 2005).

1. 2) Water-mediated chemical signalling, which has been implicated in numerous stony coral studies (Dizon & Yap, 2005; Engelhardt et al., 2023; Rinkevich & Loya, 1985a; Witzany, 2010). For instance, corals might be able to differentiate between ‘self’ and ‘non-self’ (Rinkevich, 2004) and they are surrounded by an ‘aura’ of biochemical and microbial components giving each coral colony a distinct metabolic and microbial signature (Ochsenkühn et al., 2018; Weber et al., 2019, 2022; Wegley Kelly et al., 2022), which could play a part in modulating productivity changes as observed here. 3) Acoustic communication, which is a newly arising research field that has been suggested to play a role in contact-free plant interactions (Gagliano, 2013), as it has been shown that certain terrestrial plants change their metabolism due to sound stimuli (Jung et al., 2018; Khait et al., 2019). The same has been found in the aquatic environment for macro-and microalgae (Cai et al., 2016; Frongia et al., 2020), as well as several invertebrates (Solé et al., 2023).

### Coral diversity stabilizes productivity

We found that species productivity in monoculture was a predictor for changes in productivity in polyculture. Polycultures composed of highly productive species in monoculture generally showed the highest overall productivity, indicating that these species might be less affected by the presence of other corals. Yet, these highly productive species assemblages were indeed underperforming compared to the expected productivity. Conversely, coral assemblages composed of species with low productivity in monoculture were more likely to boost their productivity in polyculture. This led to a more homogenous productivity among all stony coral assemblages, compared to larger differences between that of their monocultures. Such negative selection effects are known from grasslands (Hector et al., 2002) and marine macroalgae (Bruno et al., 2005), in which highly productive species in monoculture showed reduced productivity in polyculture and vice versa. This phenomenon is often attributed to resource partitioning in species communities (Hector et al., 2002). Our findings illustrate that negative selection effects also occur in stony corals and may contribute to robustness of productivity against changes in coral community assembly. Namely, community productivity remains relatively stable regardless of species composition, as long as genetically diverse assemblages are preserved (Clements and Hay 2021).

Interestingly, this effect was more pronounced in the conspecific assemblages compared to the multi-family assemblages. Thus, conspecific assemblages tend to enhance the productivity of less productive species and hamper the productivity of highly productivity species more than multi-family assemblages. This could be an ecological advantage for less productive species, where higher density of conspecific can increase productivity (Raymundo, 2001), which might protect them from more distantly related species (Idjadi & Karlson, 2007).

### Future directions and conclusions

With this study we add to a growing body of literature on contact-free biodiversity-productivity effects in stony corals (e.g., Clements & Hay, 2019, 2021; Dizon & Yap, 2005; McWilliam et al., 2018), yet further research linking the effects at the level of the individual organisms to interaction outcomes and the productivity balance on coral reefs is needed. Future research on communication pathways and response mechanisms within stony corals, the association with distance, and the interplay with other reef organisms will deepen our understanding on both competitive and facilitating stony coral interactions. This will enable us to evaluate changes to this interplay through climate change and other anthropogenic stressors, and aid our efforts for the successful restoration and protection of coral reefs (Cabaitan et al., 2015; Shaver & Silliman, 2017). Our research highlights the necessity of adopting a multifaceted approach to restoration of coral species and reef assemblages, recognizing that there is no one-size-fits-all solution. Certain coral species may thrive in coral nurseries through conspecific neighbours, while others may fare better within multi-family assemblages.

Our study supports the notion that corals sense the presence of neighbouring coral colonies without direct physical contact and that this results in an immediate physiological response. The direction of this response is consistent across conspecific and heterospecific neighbours, indicating a predetermined strategy to deal with other coral individuals. This process results in more homogenous productivity between diverse assemblages and potentially modulates community productivity to become more robust against shifts in species assemblages. Our study also shows that biodiversity-productivity patterns observed in terrestrial and marine plant landscapes hold for marine animals. Future studies should probe the strength of the biodiversity-productivity effects in the field, where, in comparison to terrestrial ecosystems, the within-system boundaries are less strong and physical disturbances are likely to play a more fundamental role.

## Acknowledgments

We thank Dennis Vetter for his support in implementing the mathematical theory proposed by Blomqvist (1977). This study was conducted as part of the ‘Ocean2100’ global change simulation project of the Colombian-German Center of Excellence in Marine Sciences (CEMarin) funded by the German Academic Exchange Service (DAAD). JV was supported by the German academic scholarship foundation.

## Authorship

MZ and JR conceived the ideas, MZ designed the experiment; AD, LH, AEL, and GP collected data; JV and MZ analyzed data; JV wrote the manuscript with contributions from MZ and JR. All authors read and approved the final manuscript.

## Data and Code availability

The datasets generated and analyzed during this study as well as the R scripts are available in GitHub https://github.com/JanaVetter/coral-interactions-shape-productivity.

## Supplementary information

### Supplementary Tables

**Table S1.** CITES permit numbers for the used stony coral colonies.

| Family | Species | CITES # |
| --- | --- | --- |
| Poritidae | *P. cylindrica* | 17nl241924/11 |
|  | *P. lobata* | 15-SA-000884-PD |
|  | *P. rus* | 15-SA-000885-PD |
| Acroporidae | *A. cytherea* | 19-SA-000089-PD |
|  | *A. muricata* | 14846/IV/SATS-LN/2007 |
|  | *M. digitata* | MY FSHQ/340/14 |
| Pocilloporidae | *P. damicornis* | 14846/IV/SATS-LN/2007 |
|  | *P. verrucosa* | 14846/IV/SATS-LN/2007<br>14NL214371/11<br>15-SA-000882-PD |
|  | *S. pistillata* | 15NL229192/11 |

**Table S2.**
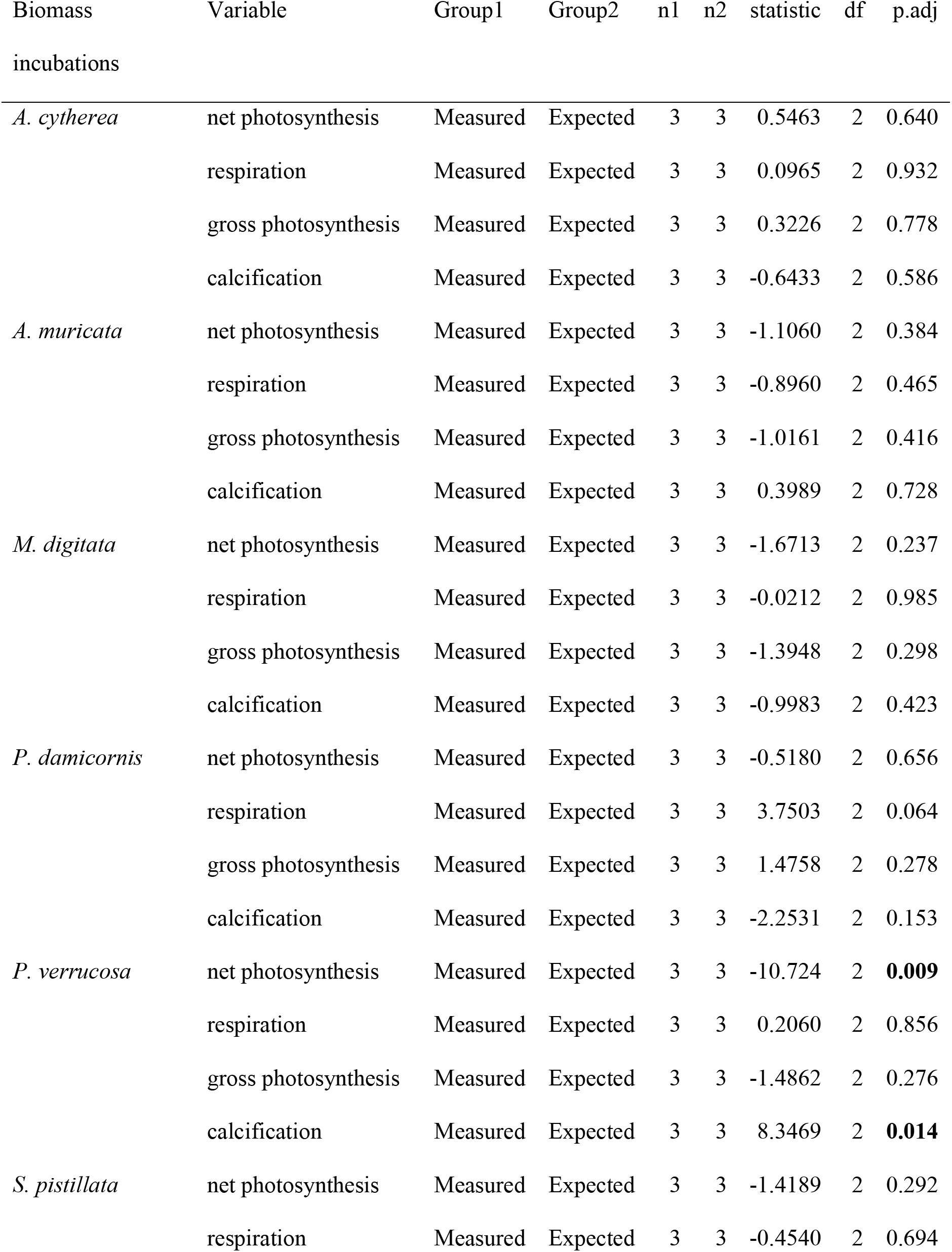
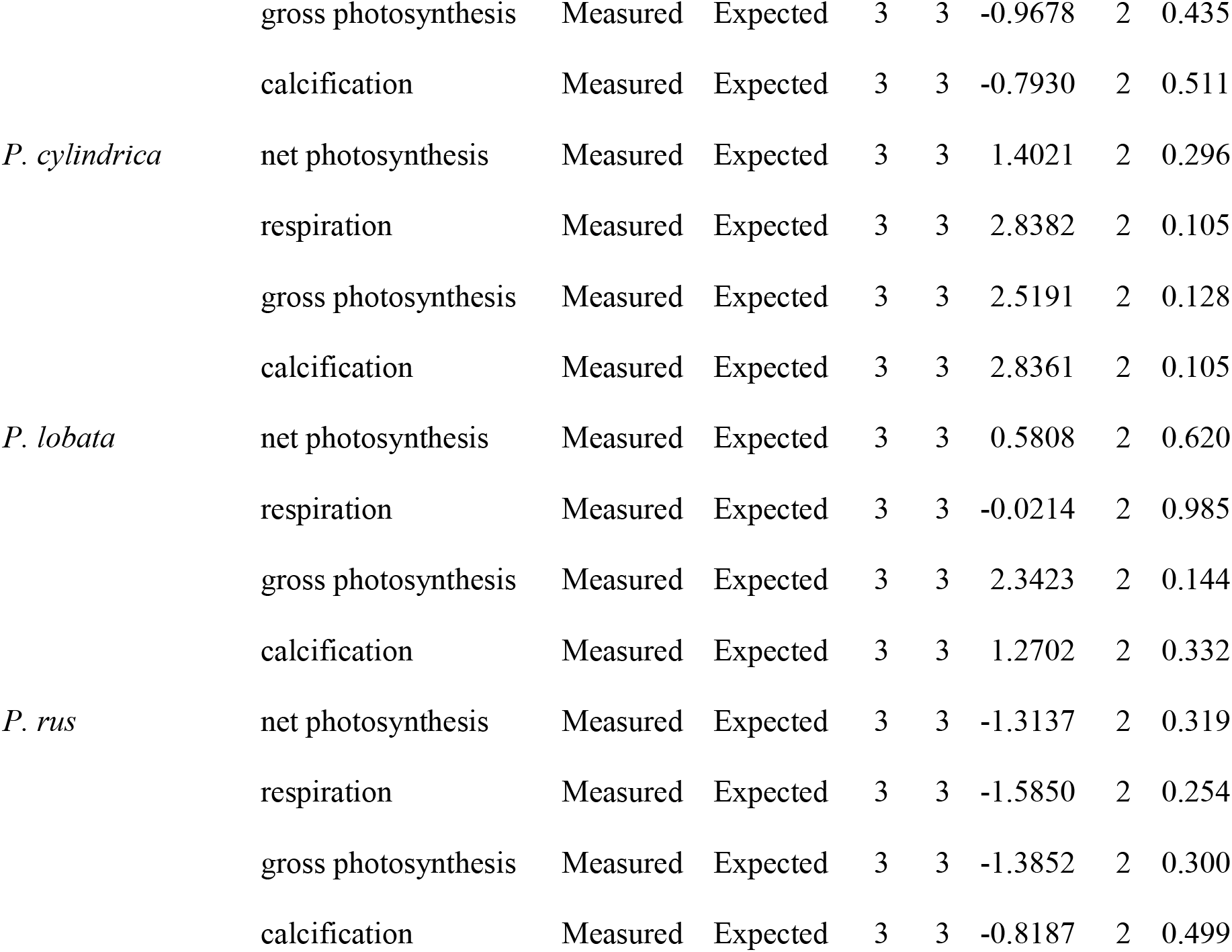
Paired t-test with Bonferroni correction comparing the mean productivity between measured biomass incubations of one species and the expected mean productivity of this species based on monoculture assemblages. Significant changes (p < 0.05) are indicated in bold.

**Table S3.**
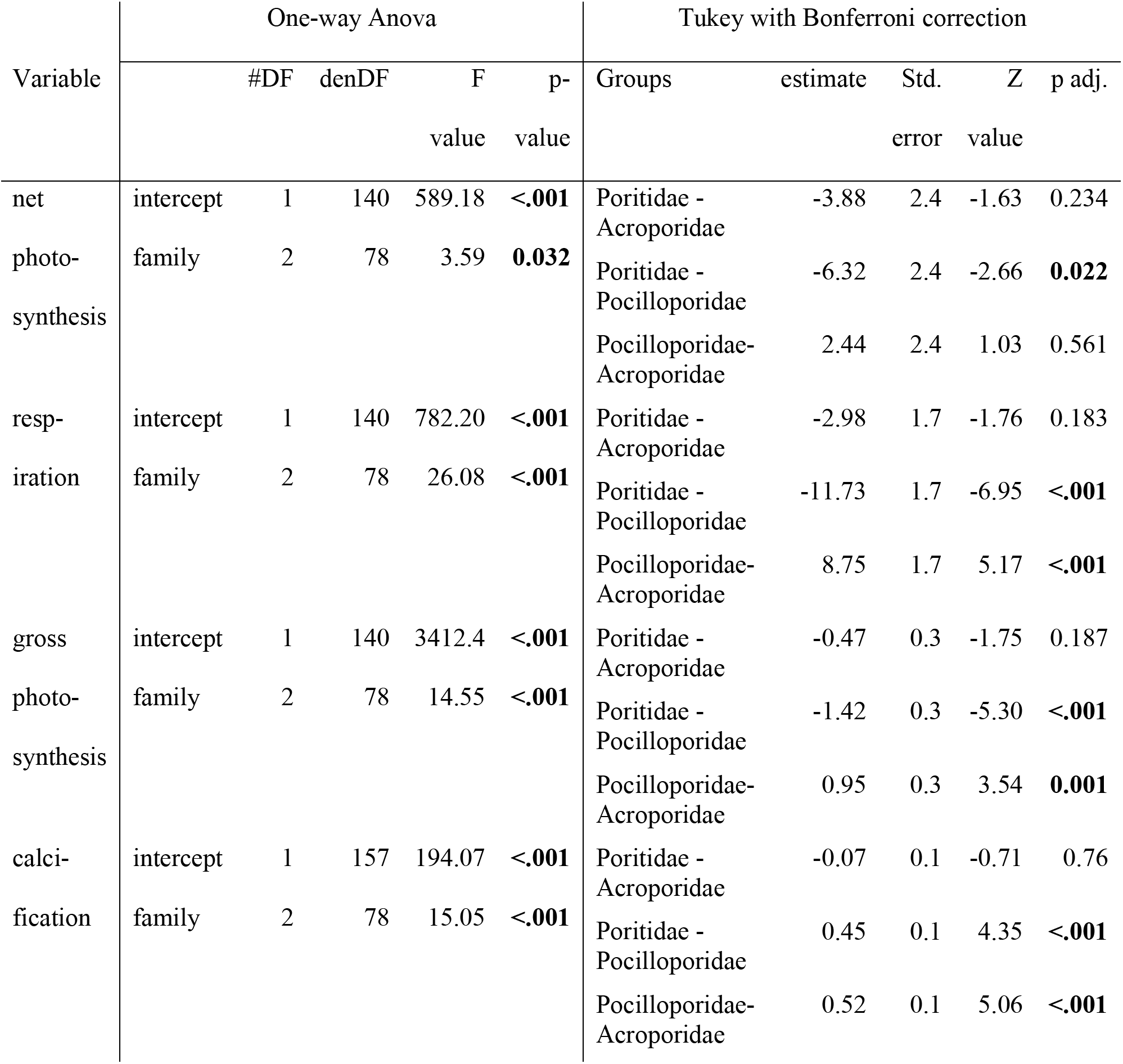
One way ANOVA of the monoculture assemblages, including coral fragment ID as random factor, comparing the mean productivity of three stony coral families. , followed by a Tukey post hoc test with Bonferroni correction (significant p < 0.05). Gross photosynthesis was square root transformed. All fragments of day 17 in runs 2 and 3 were removed from the DO measurement analysis (all monoculture incubations), because the DO_start_ values were incorrectly recorded during those runs. Additionally, five other monoculture incubations were removed from the DO calculations as outliers, as well as two outliers from the calcification calculations, due to titration problems.

**Table S4.** Productivity per species in monoculture.

| Family | Species | Productivity parameters (mean ± SD) | | | |
| --- | --- | --- | --- | --- | --- |
| | | net<br>photosynthesis<br>($\mu\text{g O}_2 \text{ cm}^{-2} \text{ h}^{-1}$) | respiration<br>($\mu\text{g O}_2 \text{ cm}^{-2} \text{ h}^{-1}$) | gross<br>photosynthesis<br>($\mu\text{g O}_2 \text{ cm}^{-2} \text{ h}^{-1}$) | calcification<br>($\mu\text{mol CaCO}_3 \text{ cm}^{-2} \text{ h}^{-1}$) |
| Poritidae | *P. cylindrica* | 13.9 ± 3.4 | 13.7 ± 4.7 | 27.6 ± 6.1 | 0.4 ± 0.4 |
| | *P. lobata* | 22.2 ± 8.0 | 16.0 ± 7.3 | 38.3 ± 13.3 | 0.6 ± 0.5 |
| | *P. rus* | 24.9 ± 10.9 | 13.7 ± 7.6 | 38.6 ± 15.6 | 0.5 ± 0.5 |
| Acroporidae | *A. cytherea* | 14.2 ± 8.6 | 15.7 ± 9.6 | 29.9 ± 16.3 | 0.1 ± 0.4 |
| | *A. muricata* | 23.2 ± 11.5 | 16.3 ± 10.0 | 39.5 ± 18.6 | 0.3 ± 0.4 |
| | *M. digitata* | 35.8 ± 6.2 | 20.6 ± 9.0 | 56.4 ± 11.2 | 0.8 ± 0.3 |
| Pocilloporidae | *P. damicornis* | 24.6 ± 7.9 | 21.5 ± 8.4 | 46.1 ± 10.3 | 0.4 ± 0.4 |
| | *P. verrucosa* | 26.4 ± 7.0 | 28.4 ± 7.0 | 54.7 ± 10.6 | 1.1 ± 0.3 |
| | *S. pistillata* | 28.7 ± 11.6 | 28.8 ± 6.5 | 57.5 ± 15.5 | 1.2 ± 0.3 |

**Table S5.**
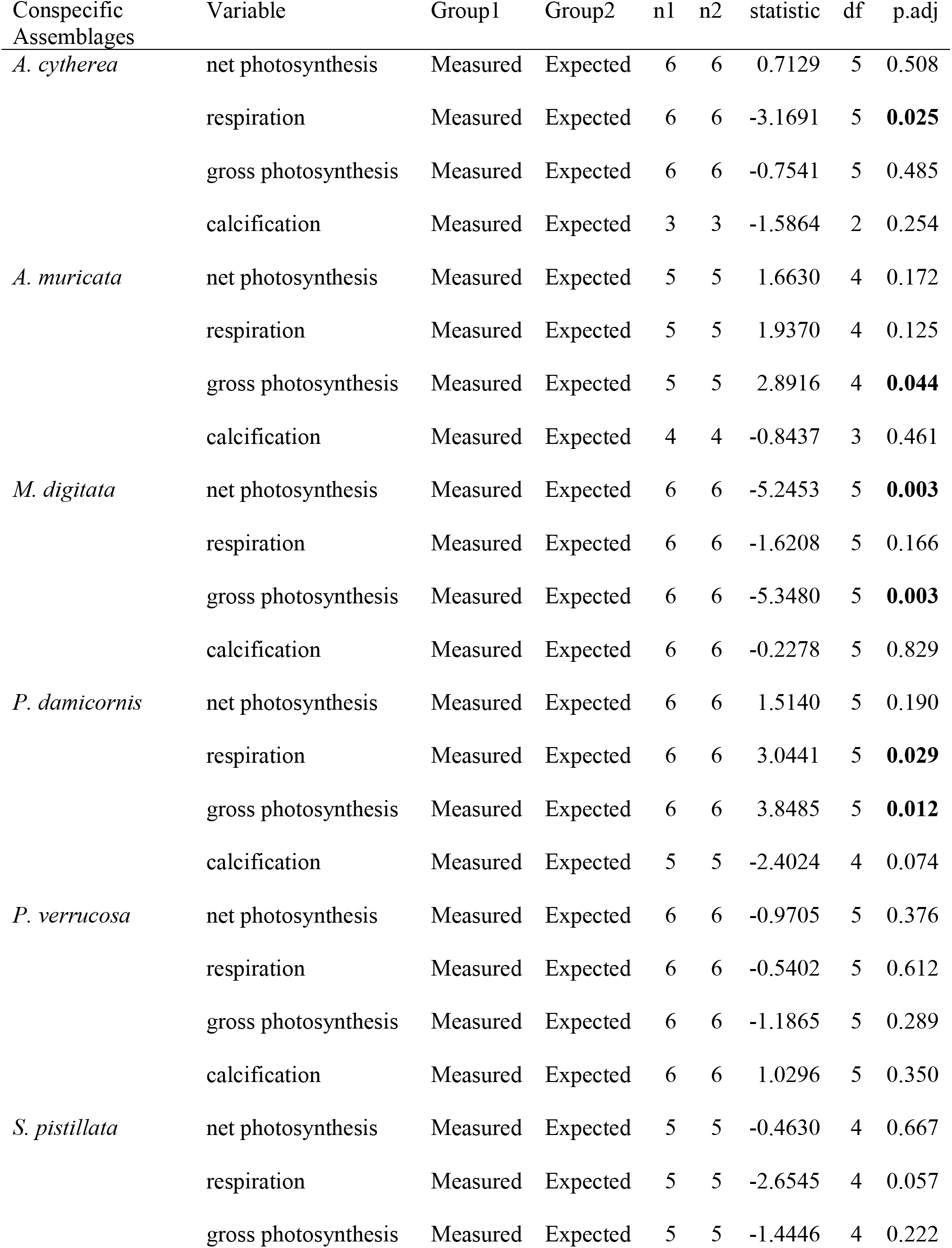
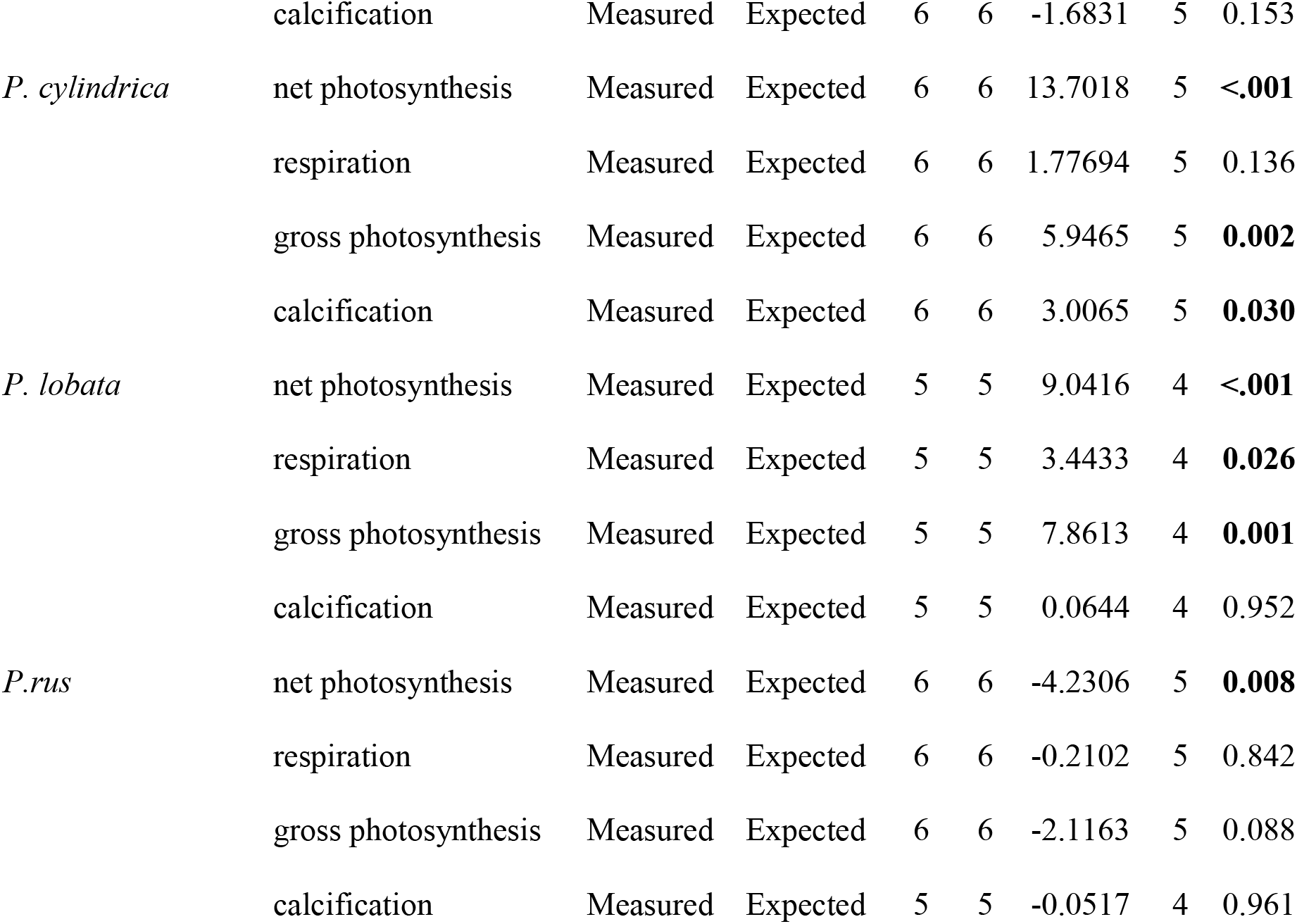
Paired t-test with Bonferroni correction comparing the measured mean productivity of conspecific incubations and the expected mean productivity of the same species based on monocultureincubations. Significant changes (p < 0.05) are indicated in bold.

**Table S6.** Paired t-test with Bonferroni correction comparing the mean productivity between measured single-family incubations and the expected mean productivity of this family. based on monoculture incubations of the species separately. Significant changes (p < 0.05) are indicated in bold.

| Single-Family Assemblages | Variable | Group1 | Group2 | n1 | n2 | statistic | df | p.adj |
| --- | --- | --- | --- | --- | --- | --- | --- | --- |
| Acroporidae | net photosynthesis | Measured | Expected | 9 | 9 | -1.4978 | 8 | 0.173 |
|  | respiration | Measured | Expected | 9 | 9 | -0.7052 | 8 | 0.501 |
|  | gross photosynthesis | Measured | Expected | 9 | 9 | -1.5125 | 8 | 0.169 |
|  | calcification | Measured | Expected | 9 | 9 | -1.1176 | 8 | 0.296 |
| Pocilloporidae | net photosynthesis | Measured | Expected | 9 | 9 | -1.0801 | 8 | 0.312 |
|  | respiration | Measured | Expected | 9 | 9 | 0.1919 | 8 | 0.853 |
|  | gross photosynthesis | Measured | Expected | 9 | 9 | -0.3223 | 8 | 0.755 |
|  | calcification | Measured | Expected | 9 | 9 | -0.0509 | 8 | 0.961 |
| Poritidae | net photosynthesis | Measured | Expected | 9 | 9 | 0.5165 | 8 | 0.619 |
|  | respiration | Measured | Expected | 9 | 9 | -1.6126 | 8 | 0.145 |
|  | gross photosynthesis | Measured | Expected | 9 | 9 | -0.5332 | 8 | 0.608 |
|  | calcification | Measured | Expected | 8 | 8 | 2.6923 | 7 | **0.031** |

**Table S7.**
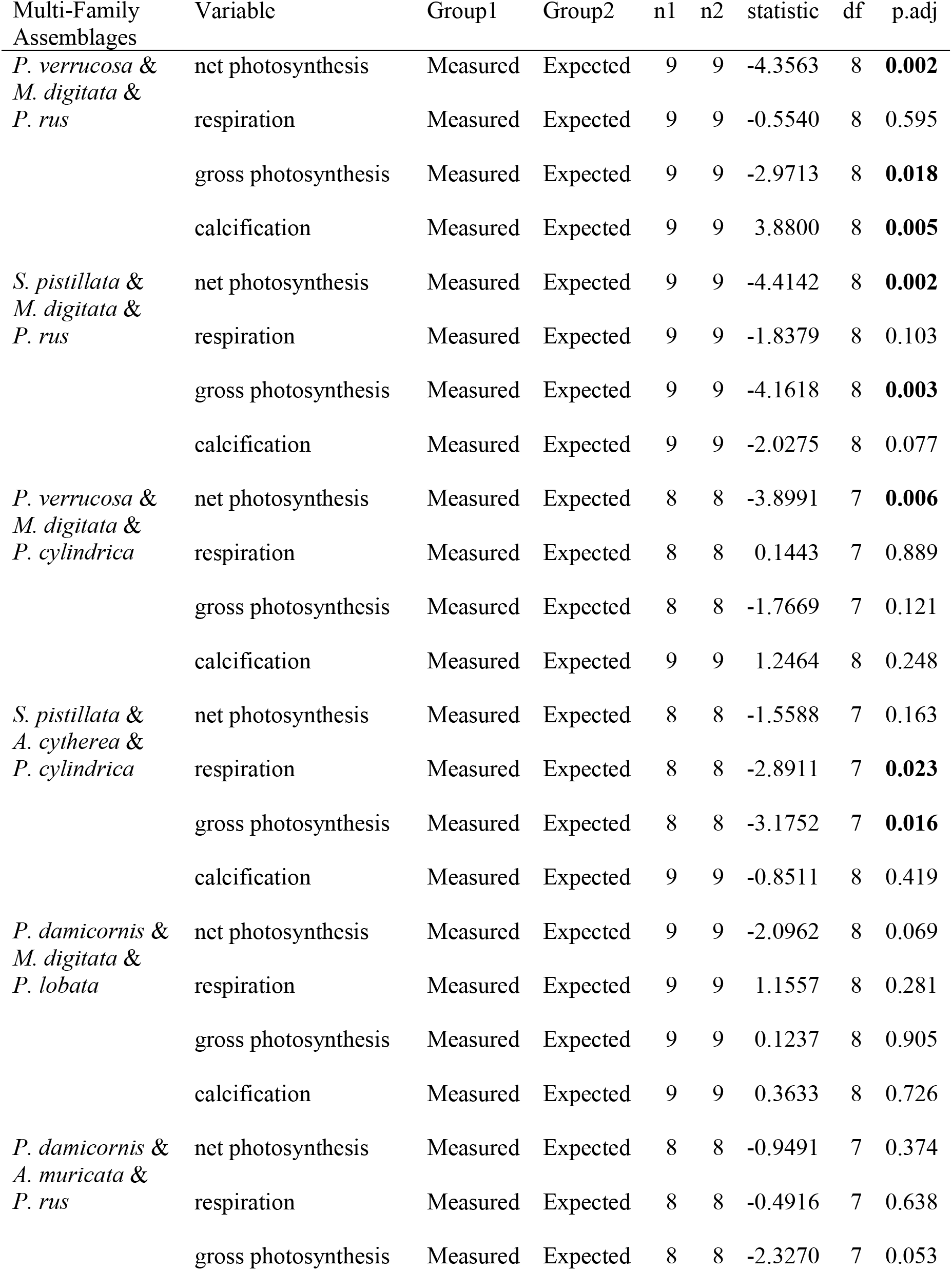
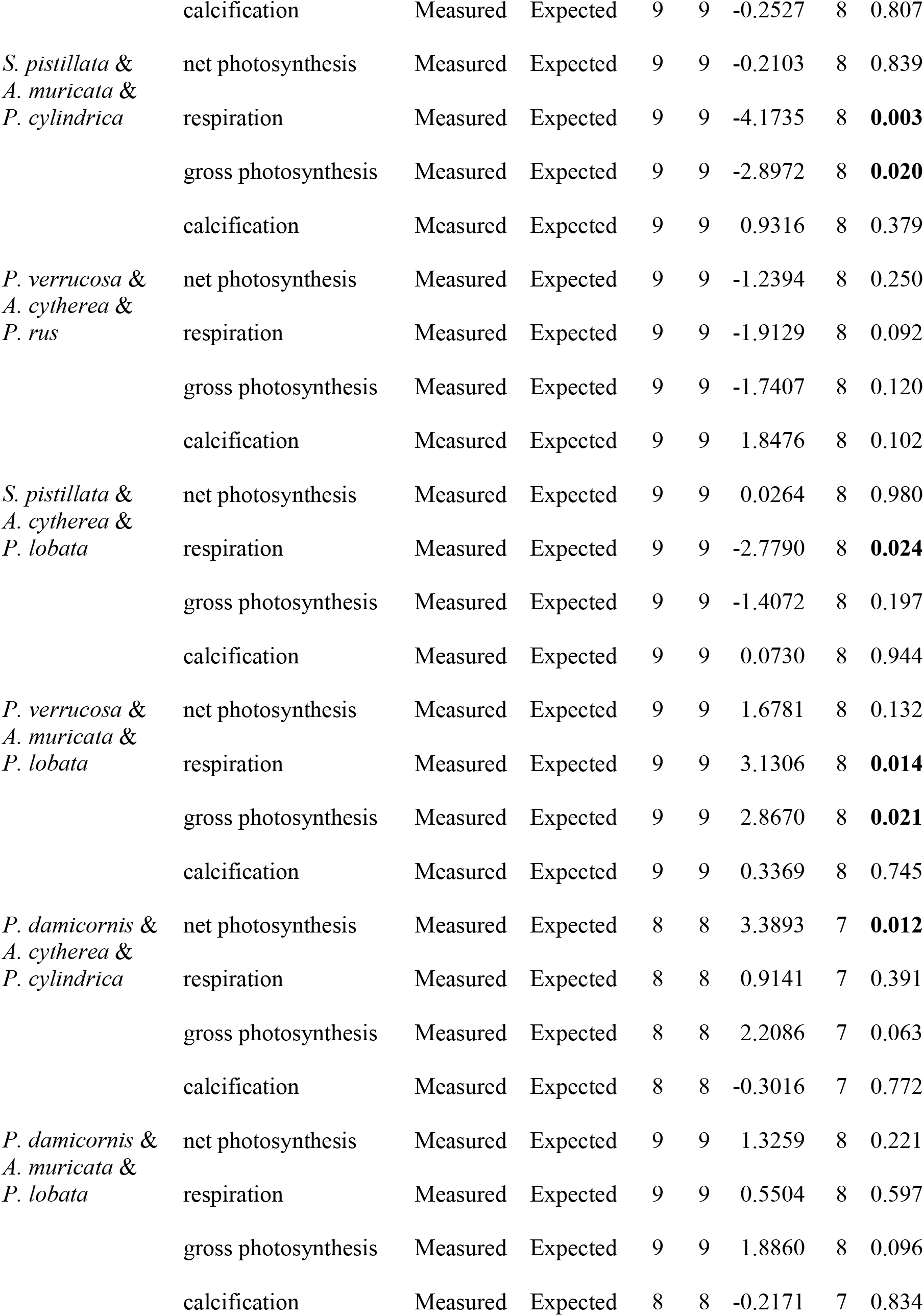
Paired t-test with Bonferroni correction comparing the mean productivity between measured multi-family incubations and the expected mean productivity. based on the sum of the monoculture incubations of the species separately (significant p < 0.05).

| Multi-Family Assemblages | Variable | Group1 | Group2 | n1 | n2 | statistic | df | p.adj |
| --- | --- | --- | --- | --- | --- | --- | --- | --- |
| *P. verrucosa* & *M. digitata* & *P. rus* | net photosynthesis | Measured | Expected | 9 | 9 | -4.3563 | 8 | **0.002** |
|  | respiration | Measured | Expected | 9 | 9 | -0.5540 | 8 | 0.595 |
|  | gross photosynthesis | Measured | Expected | 9 | 9 | -2.9713 | 8 | **0.018** |
|  | calcification | Measured | Expected | 9 | 9 | 3.8800 | 8 | **0.005** |
| *S. pistillata* & *M. digitata* & *P. rus* | net photosynthesis | Measured | Expected | 9 | 9 | -4.4142 | 8 | **0.002** |
|  | respiration | Measured | Expected | 9 | 9 | -1.8379 | 8 | 0.103 |
|  | gross photosynthesis | Measured | Expected | 9 | 9 | -4.1618 | 8 | **0.003** |
|  | calcification | Measured | Expected | 9 | 9 | -2.0275 | 8 | 0.077 |
| *P. verrucosa* & *M. digitata* & *P. cylindrica* | net photosynthesis | Measured | Expected | 8 | 8 | -3.8991 | 7 | **0.006** |
|  | respiration | Measured | Expected | 8 | 8 | 0.1443 | 7 | 0.889 |
|  | gross photosynthesis | Measured | Expected | 8 | 8 | -1.7669 | 7 | 0.121 |
|  | calcification | Measured | Expected | 9 | 9 | 1.2464 | 8 | 0.248 |
| *S. pistillata* & *A. cytherea* & *P. cylindrica* | net photosynthesis | Measured | Expected | 8 | 8 | -1.5588 | 7 | 0.163 |
|  | respiration | Measured | Expected | 8 | 8 | -2.8911 | 7 | **0.023** |
|  | gross photosynthesis | Measured | Expected | 8 | 8 | -3.1752 | 7 | **0.016** |
|  | calcification | Measured | Expected | 9 | 9 | -0.8511 | 8 | 0.419 |
| *P. damicornis* & *M. digitata* & *P. lobata* | net photosynthesis | Measured | Expected | 9 | 9 | -2.0962 | 8 | 0.069 |
|  | respiration | Measured | Expected | 9 | 9 | 1.1557 | 8 | 0.281 |
|  | gross photosynthesis | Measured | Expected | 9 | 9 | 0.1237 | 8 | 0.905 |
|  | calcification | Measured | Expected | 9 | 9 | 0.3633 | 8 | 0.726 |
| *P. damicornis* & *A. muricata* & *P. rus* | net photosynthesis | Measured | Expected | 8 | 8 | -0.9491 | 7 | 0.374 |
|  | respiration | Measured | Expected | 8 | 8 | -0.4916 | 7 | 0.638 |
|  | gross photosynthesis | Measured | Expected | 8 | 8 | -2.3270 | 7 | 0.053 |
|  | calcification | Measured | Expected | 9 | 9 | -0.2527 | 8 | 0.807 |
| *S. pistillata* &<br>*A. muricata* &<br>*P. cylindrica* | net photosynthesis | Measured | Expected | 9 | 9 | -0.2103 | 8 | 0.839 |
|  | respiration | Measured | Expected | 9 | 9 | -4.1735 | 8 | **0.003** |
|  | gross photosynthesis | Measured | Expected | 9 | 9 | -2.8972 | 8 | **0.020** |
|  | calcification | Measured | Expected | 9 | 9 | 0.9316 | 8 | 0.379 |
| *P. verrucosa* &<br>*A. cytherea* &<br>*P. rus* | net photosynthesis | Measured | Expected | 9 | 9 | -1.2394 | 8 | 0.250 |
|  | respiration | Measured | Expected | 9 | 9 | -1.9129 | 8 | 0.092 |
|  | gross photosynthesis | Measured | Expected | 9 | 9 | -1.7407 | 8 | 0.120 |
|  | calcification | Measured | Expected | 9 | 9 | 1.8476 | 8 | 0.102 |
| *S. pistillata* &<br>*A. cytherea* &<br>*P. lobata* | net photosynthesis | Measured | Expected | 9 | 9 | 0.0264 | 8 | 0.980 |
|  | respiration | Measured | Expected | 9 | 9 | -2.7790 | 8 | **0.024** |
|  | gross photosynthesis | Measured | Expected | 9 | 9 | -1.4072 | 8 | 0.197 |
|  | calcification | Measured | Expected | 9 | 9 | 0.0730 | 8 | 0.944 |
| *P. verrucosa* &<br>*A. muricata* &<br>*P. lobata* | net photosynthesis | Measured | Expected | 9 | 9 | 1.6781 | 8 | 0.132 |
|  | respiration | Measured | Expected | 9 | 9 | 3.1306 | 8 | **0.014** |
|  | gross photosynthesis | Measured | Expected | 9 | 9 | 2.8670 | 8 | **0.021** |
|  | calcification | Measured | Expected | 9 | 9 | 0.3369 | 8 | 0.745 |
| *P. damicornis* &<br>*A. cytherea* &<br>*P. cylindrica* | net photosynthesis | Measured | Expected | 8 | 8 | 3.3893 | 7 | **0.012** |
|  | respiration | Measured | Expected | 8 | 8 | 0.9141 | 7 | 0.391 |
|  | gross photosynthesis | Measured | Expected | 8 | 8 | 2.2086 | 7 | 0.063 |
|  | calcification | Measured | Expected | 8 | 8 | -0.3016 | 7 | 0.772 |
| *P. damicornis* &<br>*A. muricata* &<br>*P. lobata* | net photosynthesis | Measured | Expected | 9 | 9 | 1.3259 | 8 | 0.221 |
|  | respiration | Measured | Expected | 9 | 9 | 0.5504 | 8 | 0.597 |
|  | gross photosynthesis | Measured | Expected | 9 | 9 | 1.8860 | 8 | 0.096 |
|  | calcification | Measured | Expected | 8 | 8 | -0.2171 | 7 | 0.834 |

**Table S8.** Wilcoxon rank-sum test output between the change from expected productivity (%) of conspecific and multi-family assemblages. . Significant changes (p < 0.05) are indicated in bold.

| Variable | Group1 | Group2 | W | p |
| --- | --- | --- | --- | --- |
| Net photosynthesis change from expected | conspecific | Multi-Family | 3697 | **< 0.001** |
| Respiration change from expected | conspecific | Multi-Family | 3026 | 0.155 |
| Gross photosynthesis change from expected | conspecific | Multi-Family | 3410 | **0.004** |
| Calcification change from expected | conspecific | Multi-Family | 2600 | 0.517 |

**Table S9.**
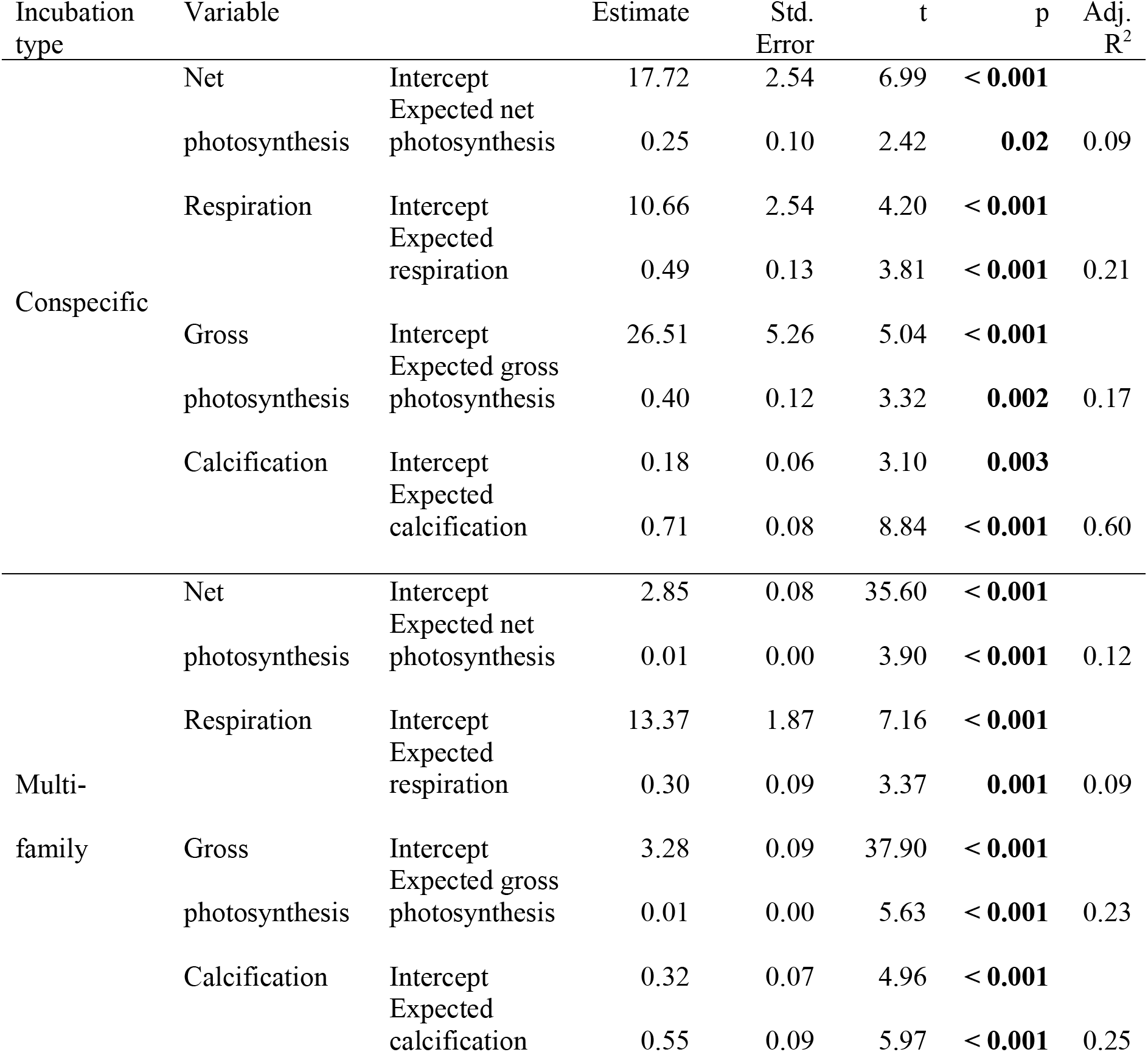
Linear regression output to compare between measured productivity and expected productivity of the conspecific (n = 51) and multi-family (n = 104) assemblages. . Measured net photosynthesis and measured gross photosynthesis of the multi-family incubations was log transformed.

**Table S10.** Pearson correlation output for comparison between the change from expected (measured - expected) and the expected productivity of conspecific (n = 51) and multi-family (n = 104) assemblages. R was adjusted according to Blomqvist (1977) and corrected p-values were obtained via Fisher’s z-transformation.

| Incubation type | Variable | Original R | adj. R | 95% Conf. | adj. p |
| --- | --- | --- | --- | --- | --- |
| Conspecific | Net photosynthesis | -0.73 | -0.65 | -0.79 | **<0.001** |
|  |  |  |  | -0.46 |  |
|  | Respiration | -0.50 | -0.25 | -0.49 | 0.08 |
|  |  |  |  | 0.03 |  |
|  | Gross photosynthesis | -0.59 | -0.47 | -0.66 | **< 0.001** |
|  |  |  |  | -0.23 |  |
|  | Calcification | -0.45 | -0.21 | -0.46 | 0.14 |
|  |  |  |  | 0.07 |  |
| Multi-family | Net photosynthesis | -0.69 | -0.22 | -0.40 | **0.02** |
|  |  |  |  | -0.03 |  |
|  | Respiration | -0.62 | -0.29 | -0.46 | **0.002** |
|  |  |  |  | -0.11 |  |
|  | Gross photosynthesis | -0.56 | -0.34 | -0.50 | **< 0.001** |
|  |  |  |  | -0.16 |  |
|  | Calcification | -0.43 | 0.12 | -0.07 | 0.20 |
|  |  |  |  | 0.31 |  |

### Supplementary Figures

**Figure S1.**
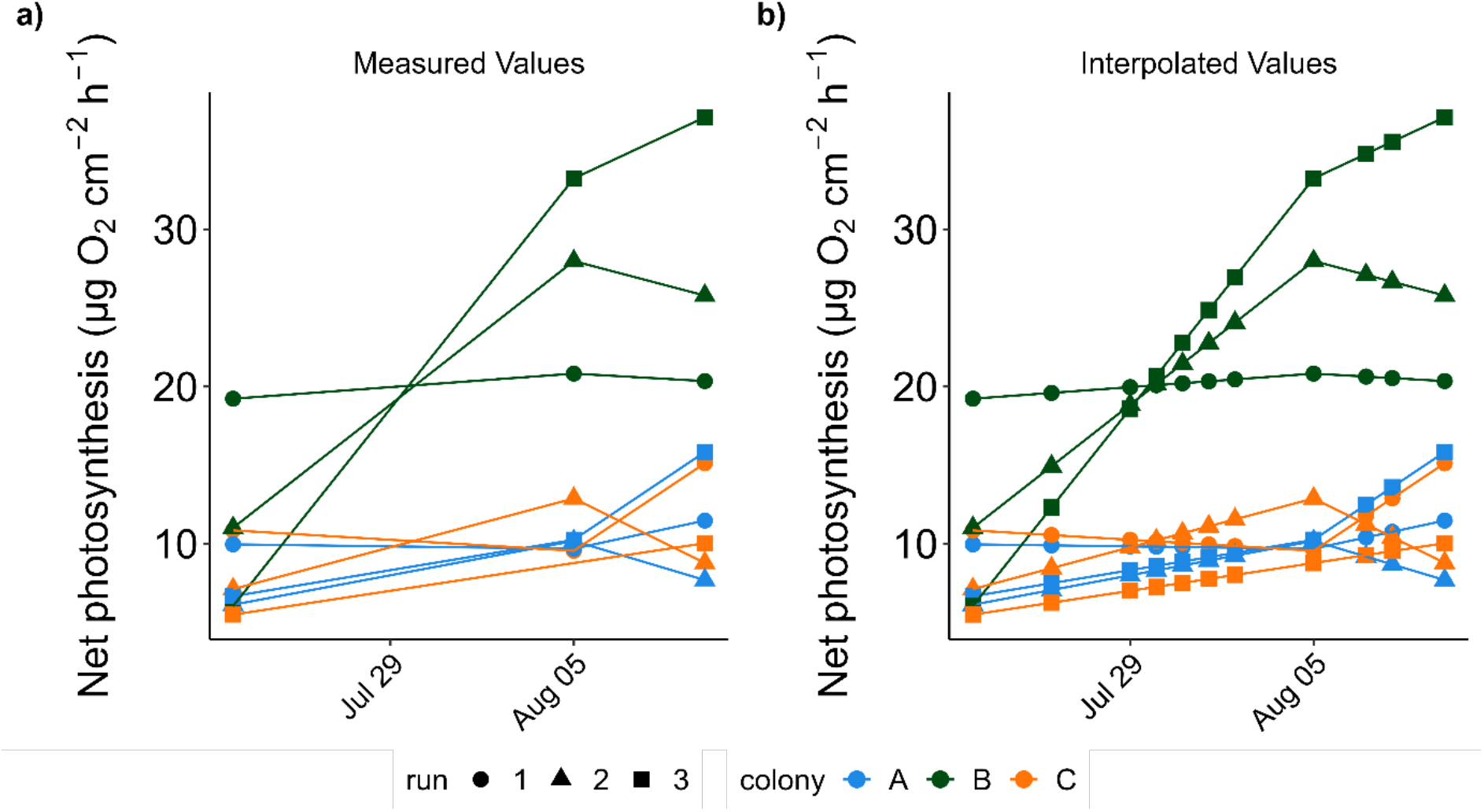
Example of productivity value interpolation between monocultures. This example shows the interpolation of *A. cytherea* net photosynthesis values, which were later used to calculate the expected values of polyculture incubations in which *A. cytherea* was present. A linear relationship was assumed between two measured points. Interpolated values were only calculated for the necessary dates (in which a polyculture incubation with the species was conducted) and the values were normalized to 1 cm^2^ coral tissue surface area. These were matched with the polyculture incubation day, colonies, and run, and multiplied by the matching fragments surface area, to account for the unique contribution by different sized fragments, to the combined polyculture productivity. These combined productivity values were then divided by the sum of the surface area of all the fragments of the incubation, so that values are expressed per cm^2^, for comparability among all incubations.

**Figure S2.**
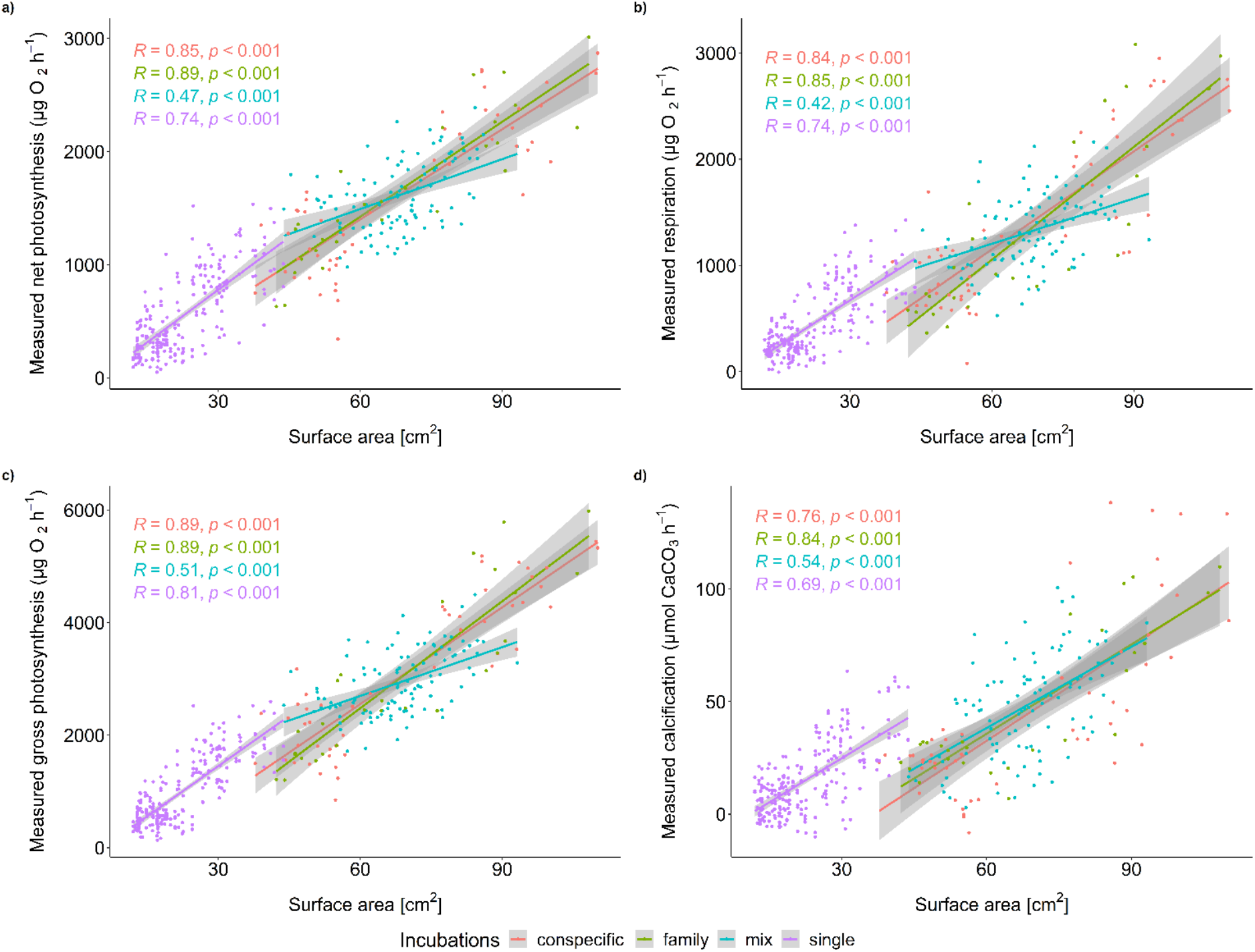
Pearson correlation between surface area and productivity parameters: net photosynthesis (a), respiration (b), gross photosynthesis (c) and calcification (d).

**Figure S3.**
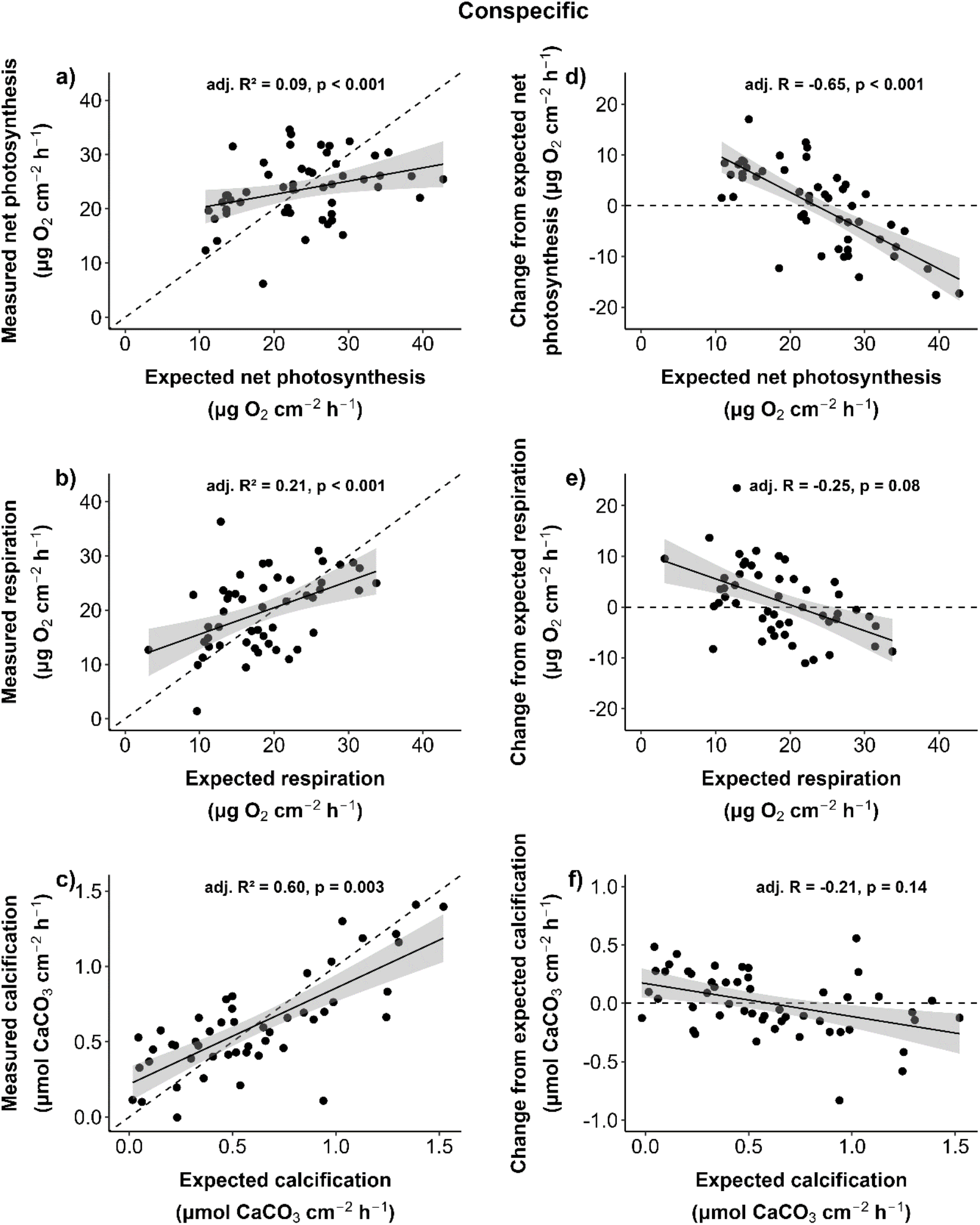
Relationship between productivity of corals in monoculture and conspecific assemblages. Data shown as the relationship between the measured productivity of conspecific assemblages and their expected productivity, displayed with adjusted R^2^ and p-values from a linear regression model (a-c). The productivity change of conspecific assemblages (measured - expected) and their expected productivity are shown with corrected R and p-value following Blomqvist (1977, d-f). Dashed lines indicate the 1:1 productivity (no change). Points above the dashed lines are assemblages which were more productive in conspecific assemblages (n = 51).

**Figure S4.**
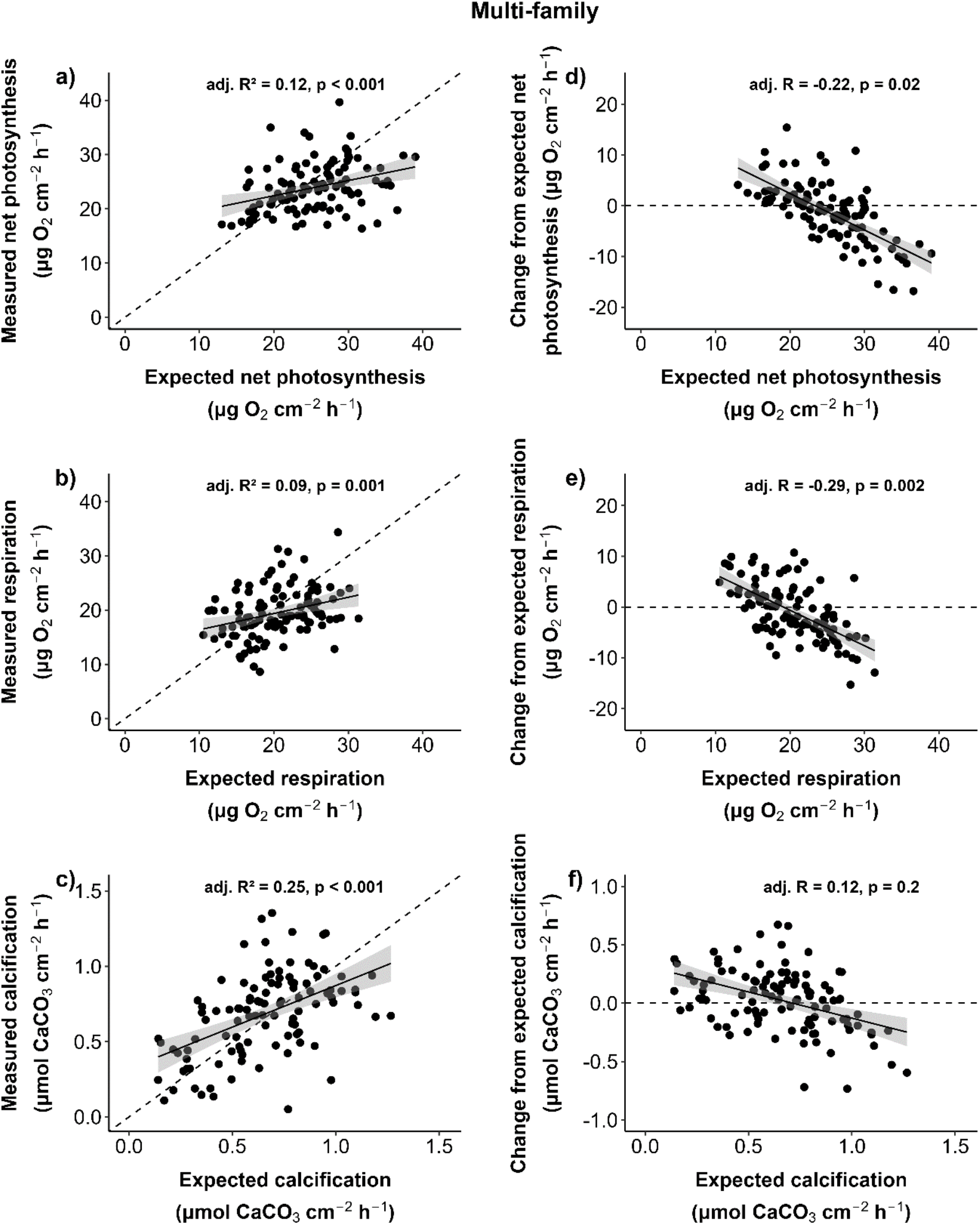
Relationship between productivity of corals in monoculture and multi-family assemblages. Data shown as the relationship between the measured productivity of multi-family assemblages and their expected productivity, displayed with adjusted R^2^ and p-values from a linear regression model (a-c). The productivity change of multi-family assemblages (measured - expected) and their expected productivity are shown with corrected R and p-value following Blomqvist (1977, d-f). Dashed lines indicate the 1:1 productivity (no change). Points above the dashed lines are assemblages which were more productive in multi-family assemblages (n = 104).

